# ROS-dependent palmitoylation is an obligate licensing modification for GSDMD pore formation

**DOI:** 10.1101/2023.03.07.531538

**Authors:** Gang Du, Liam B. Healy, Liron David, Caitlin Walker, Pietro Fontana, Ying Dong, Pascal Devant, Robbins Puthenveetil, Scott B. Ficarro, Anirban Banerjee, Jonathan C. Kagan, Judy Lieberman, Hao Wu

## Abstract

Gasdermin D (GSDMD) is the common effector for cytokine secretion and pyroptosis downstream of inflammasome activation by forming large transmembrane pores upon cleavage by inflammatory caspases. Here we report the surprising finding that GSDMD cleavage is not sufficient for its pore formation. Instead, GSDMD is lipidated by S-palmitoylation at Cys191 upon inflammasome activation, and only palmitoylated GSDMD N-terminal domain (GSDMD-NT) is capable of membrane translocation and pore formation, suggesting that palmitoylation licenses GSDMD activation. Treatment by the palmitoylation inhibitor 2-bromopalmitate and alanine mutation of Cys191 abrogate GSDMD membrane localization, cytokine secretion, and cell death, without affecting GSDMD cleavage. Because palmitoylation is formed by a reversible thioester bond sensitive to free thiols, we tested if GSDMD palmitoylation is regulated by cellular redox state. Lipopolysaccharide (LPS) mildly and LPS plus the NLRP3 inflammasome activator nigericin markedly elevate reactive oxygen species (ROS) and GSDMD palmitoylation, suggesting that these two processes are coupled. Manipulation of cellular ROS by its activators and quenchers augment and abolish, respectively, GSDMD palmitoylation, GSDMD pore formation and cell death. We discover that zDHHC5 and zDHHC9 are the major palmitoyl transferases that mediate GSDMD palmitoylation, and when cleaved, recombinant and partly palmitoylated GSDMD is 10-fold more active in pore formation than bacterially expressed, unpalmitoylated GSDMD, evidenced by liposome leakage assay. Finally, other GSDM family members are also palmitoylated, suggesting that ROS stress and palmitoylation may be a general switch for the activation of this pore-forming family.

**One-Sentence Summary:** GSDMD palmitoylation is induced by ROS and required for pore formation

---

Since the discovery of Gasdermin D (GSDMD) as a substrate of inflammatory caspases (1, 4, 5, and 11) that form transmembrane pores upon cleavage (*1-9*), the GSDM family proteins have been implicated as key players in host defense and homeostatic control by mediating the release of pro-inflammatory cytokines and alarmins, and executing the programmed inflammatory cell death known as pyroptosis (*9, 10*). In addition to activation of GSDMD by caspase cleavage, all GSDMs are now known to be activated by specific host and microbial proteases under various physiological contexts, from apoptotic caspase-3 and -8, granzymes delivered to target cells by cytotoxic lymphocytes, neutrophil granule proteases released to the cytosol, to *Streptococcal* proteases (*9, 11*). These data established the paradigm that GSDMs are activated by proteolytic processing in between the pore-forming N-terminal domain (NT) and the autoinhibitory C-terminal domain (CT). Crystal structures of autoinhibited, full-length monomeric GSDMs (*4, 12*) and cryo-EM structures of GSDM-NTs in transmembrane pore form or oligomeric prepore form before membrane insertion (*7, 8*) further consolidated this paradigm.

Protein S-palmitoylation is a powerful post-translational modification that adds a long lipid chain to Cys residues in proteins via a thioester bond to regulate localization, trafficking, and other important cellular processes (*13, 14*). It is also the only reversible post-translational lipid modification, which could serve to control the signal transduction of both soluble and integral membrane proteins such as Ras, and STING, respectively (*15-17*). Dysregulation of protein palmitoylation could lead to important human diseases from neurological disorders to cancer.

Reactive oxygen species (ROS) are composed of a number of heterogeneous chemical entities that induce oxidative stress (*18*) and have long been recognized as an important factor in NLRP3 activation and inflammasome formation (*19-21*). Intriguing, recent studies also implicated a role of ROS in GSDMD-NT pore formation independent of upstream processes (*22, 23*), but the mechanism of this role remained unclear.

Here we found that GSDMD is S-palmitoylated at Cys191 in human GSDMD within its NT (Cys192 in mouse GSDMD), a residue shown to be crucial for GSDMD-NT-induced cell death in cells (*3*). Importantly, this palmitoylation is ROS-dependent and regulated, which differs from previously reported GSDME palmitoylation at its CT (*24*) or autocatalytic bacterial GSDM palmitoylation near its N-terminus (*25*). Our findings explain the ROS dependence of GSDMD-NT pore formation, revealing a specific effect by palmitoylation downstream of the rather non-specific ROS, and reveals an obligate requirement for palmitoylation in GSDMD-NT pore formation in addition to its cleavage.

## GSDMD is palmitoylated and only the membrane has palmitoylated GSDMD-NT

We observed that the C191A mutant of GSDMD-NT is impaired in oligomerization and cell death induction (*3*). We hypothesized that C191 might be involved in a disulfide bond formation in the GSDMD-NT pore; however, there are no such bonds in either the full-length autoinhibited form (PDB ID: 6N9O) (*12*), or the membrane-inserted pore form (PDB ID: 6VFE) (*8*) of GSDMD, and the distance between C191 residues in the GSDMD-NT pore is ∼20 Å, which is too long for a disulfide bond to bridge (**Fig. 1A, fig. S1A**). In addition, the structure of the GSDMD-NT pore does not suggest a mechanism to explain the defectiveness of the C191A mutant. To address whether there might be a cell-specific mechanism for the impairment of pore formation by C191A, we compared pore formation activity by bacterially expressed WT GSDMD and the C191A mutant in a liposome leakage assay upon cleavage to generate the active GSDMD-NT. Kinetic monitoring for the leakage of encapsulated Tb^3+^ ions in the liposomes revealed similar activity of WT and C191A (**fig. S1B**), suggesting the existence of a cell-specific mechanism.

**Fig. 1.**
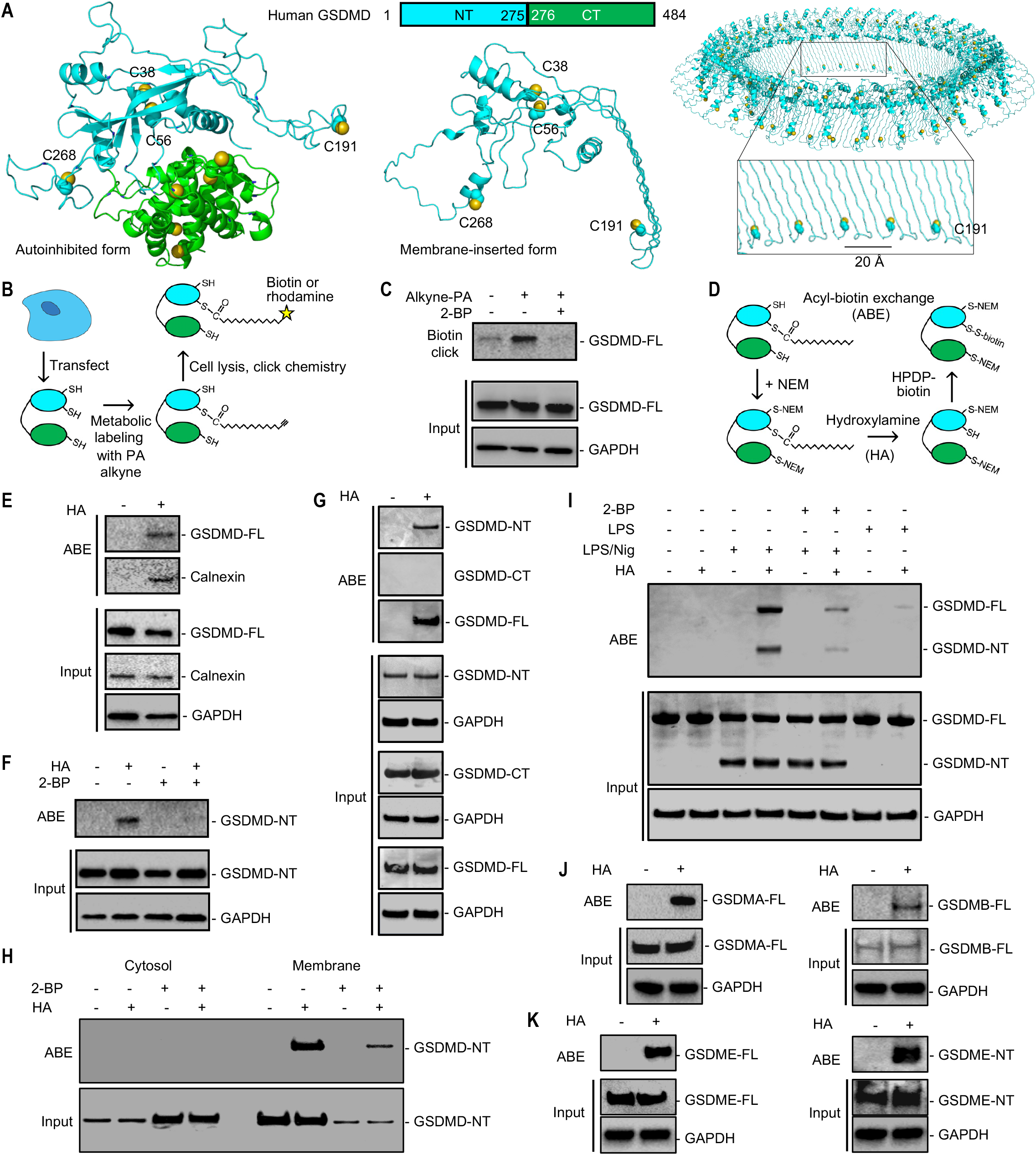
GSDMD is highly palmitoylated, only the membrane has palmitoylated GSDMD, and other GSDMs are also palmitoylated. (**A**) Domain organization and ribbon diagrams of autoinhibited full-length (FL) GSDMD (modeled after PDB 6N9O (*12*), left), membrane-inserted GSDMD-NT monomer (middle) and the GSDMD-NT pore oligomer modeled after PDB 6VFE (*8*) (right). (**B**) A schematic of metabolic labeling and click chemistry for detecting protein acylation. (**C**) GSDMD-FL palmitoylation when overexpressed in HEK293T cells, detected by alkyne-palmitic acid (PA) labeling, biotin-azide click, and biotin pulldown, and its inhibition by the general palmitoylation inhibitor 2-BP. GAPDH acts as the loading control. (**D**) A schematic of acyl-biotin exchange (ABE) for detecting protein acylation. Free thiols on proteins are blocked by N-ethylmaleimide (NEM), followed by treatment with hydroxylamine (HA) to break acylated thiols to generate free thiols. Free thiols are biotinylated by thiol-reactive biotin (HPDP-biotin), biotinylated proteins are pulled down by streptavidin beads, and specific proteins are detected by western blots. (**E**) GSDMD-FL palmitoylation when overexpressed in HEK293T cells detected by ABE, with the known palmitoylated protein Calnexin as a positive control. (**F**) GSDMD-NT palmitoylation detected by ABE and its inhibition by 2-BP. (**G**) Palmitoylation of GSDMD-FL and GSDMD-NT, but not GSDMD-CT, detected by ABE. (**H**) Palmitoylated GSDMD-NT in the membrane, but not the cytosolic fraction, detected by ABE following cell lysate fractionation. (**I**) Palmitoylation of GSDMD-FL and GSDMD-NT in THP-1 cells upon stimulation or not by LPS or LPS plus nigericin. 2-BP inhibited GSDMD palmitoylation without affecting its processing. (**J, K**) Palmitoylation of GSDMA-FL and GSDMB-FL (J), and GSDME-FL and GSDME-NT (K), detected by ABE. All GSDMs were overexpressed in HEK293T cells.

Because GSDMD functions in membranes and cysteines can be reversibly palmitoylated in cells, we suspected that GSDMD may be S-palmitoylated at C191, which may explain why C191A is defective in pore formation. To determine experimentally if GSDMD is palmitoylated, we first used alkyne-azide click chemistry with metabolic labeling (*26*) (**Fig. 1B**). HEK293T cells were transfected with full-length human GSDMD and treated with alkyne palmitic acid (also known as hexadecynoic acid, or 15-HDYA), a clickable analog of palmitic acid, to metabolic label sites of palmitoylation. Upon cell lysis, azido-biotin was added to chemically attach biotin to palmitoylated proteins. A pulldown using streptavidin-coated beads followed by immunoblotting against GSDMD revealed robust GSDMD palmitoylation (**Fig. 1C**). By contrast, palmitoylated GSDMD was nearly absent either without treatment by alkyne palmitic acid or with additional treatment by 2-bromopalmitate (2-BP), a non-metabolizable palmitate analog and a general palmitoylation inhibitor that binds to the active site of palmitoyl acyl-transferases. The GSDMD expression levels were equal among the different conditions, as was the GAPDH control.

Because different forms of lipidation often work together to promote dynamic and regulated localization to specific membranes (*15*), we also ran predictions on other forms of lipidation using GPS-Lipid (*27*), which detected a non-consensus N-myristoylation site, with potential N-myristoylation at residue G2. However, click chemistry after metabolic labeling (*26*) using alkyne myristic acid, a clickable analog of myristic acid, did not result in detectable N-myristoylation (**fig. S1C**), while the control experiment detected robust palmitoylation (**fig. S1D**).

To confirm GSDMD palmitoylation, we used an alternative method, acyl-biotin exchange (ABE) (*26*) (**Fig. 1D**), in which cell lysates were first treated with the alkylation reagent N-ethylmaleimide (NEM) to block unmodified Cys residues. This was then followed by treatment with hydroxylamine (HA, to remove palmitoylation) or NaCl (negative control), and addition of a pyridyldithiol-biotin compound (biotin-HPDP) to label free Cys residues (*26*). Biotinylated proteins were then pulled down using streptavidin beads, and immunoblotted to detect GSDMD (**Fig. 1E**). With, but not without, HA treatment, biotinylated GSDMD was detected, which further demonstrated that GSDMD is palmitoylated. A protein known to undergo palmitoylation, calnexin, was used as a positive control in the same experiment. Using ABE, we performed a similar experiment on GSDMD-NT and GSDMD-CT. We found that GSDMD-NT was palmitoylated, and this palmitoylation was blocked by treatment with 2-BP, while GSDMD-CT was not palmitoylated (**Fig. 1F, G**).

To examine the localization of GSDMD-NT with or without palmitoylation, we fractionated lysates from GSDMD-NT expressing HEK293T cells into cytosolic and membrane fractions. ABE in each of the fractions detected palmitoylated GSDMD-NT only in the membrane fraction, but none in the cytosolic fraction, and 2-BP reduced GSDMD-NT palmitoylation (**Fig. 1H**). In this experiment, the percentage of GSDMD-NT palmitoylation estimated by the ABE band intensity versus the total input band intensity (2-fold diluted) is ∼35%. With 2-BP treatment, palmitoylation was reduced to an estimated ∼8%. These data suggest that palmitoylation is linked to GSDMD-NT poring activity.

To investigate GSDMD palmitoylation during inflammasome activation, we performed ABE on differentiated THP-1 cells, a human monocytic cell line that expresses endogenously inflammasome pathway proteins. GSDMD palmitoylation was not detected with HA treatment alone, and priming by lipopolysaccharides (LPS) slightly increased biotinylated GSDMD. By contrast, LPS priming followed by nigericin treatment to activate the NLRP3 inflammasome markedly enhanced GSDMD palmitoylation, and 2-BP significantly blocked GSDMD palmitoylation (**Fig. 1I**). Thus, inflammasome activation is associated with heightened GSDMD palmitoylation. Of note, GSDMD cleavage was not affected by 2-BP treatments (**Fig. 1I**); thus inhibition of palmitoylation did not affect the upstream steps in NLRP3 activation.

### Other GSDMs are also palmitoylated

The high degree of GSDMD palmitoylation and the universal function of GSDMs in membrane pore formation promoted us to ask if other GSDMs might also be palmitoylated. To address this question, we transfected all other GSDMs (A, B, C, E) in HEK293T cells, and performed ABE. These data showed that GSDMA, B, and E were all palmitoylated (**Fig. 1J, K**); we did not detect GSDMC expression. Specifically, palmitoylation was detected at GSDME-NT, which is different from the previously reported GSDME-CT palmitoylation **(*24*)**. Since CT is not part of the transmembrane pore, we propose that the observed inhibition of chemotherapy-induced pyroptosis by 2-BP (*24*) is from its effect on GSDME-NT, and the GSDME-dependent anticancer effect of Erianin is through suppression of a depalmitoylase that enzymatically removes the palmitate (*28*). These data suggest that palmitoylation, possibly all at GSDM-NTs, is an obligate requirement for GSDM pore formation.

### GSDMD is palmitoylated only at Cys191, which is required for membrane localization and cell death by GSDMD-NT

There are four Cys residues at the GSDMD-NT (**fig. S1A**), To determine which Cys is palmitoylated, we generated GSDMD-NT constructs with single alanine mutations as well as the I104N mutation (I105N mutation in mouse GSDMD), which was shown to reduce GSDMD function in the NT form (*1, 29*). We transfected WT and mutant GSDMD-NT in HEK293T cells and performed ABE to examine their palmitoylation. All mutants behaved as the WT except the C191A mutant for which no palmitoylation could be detected (**Fig. 2A**). This strong selectivity for C191 is notable, especially in comparison with oxidation (*23*). The C191A mutant, but not other Cys mutants, exhibited reduced cell death shown by propidium iodide (PI) staining (**Fig. 2B, fig. S2A**), lactate dehydrogenase (LDH) release (**Fig. 2C**), and the ATP luminescent cell viability assay (**Fig. 2D**). Immunofluorescence further showed that while WT GSDMD-NT localized primarily to the cell surface, C191A exhibited largely cytosolic staining (**Fig. 2E**).

**Fig. 2.**
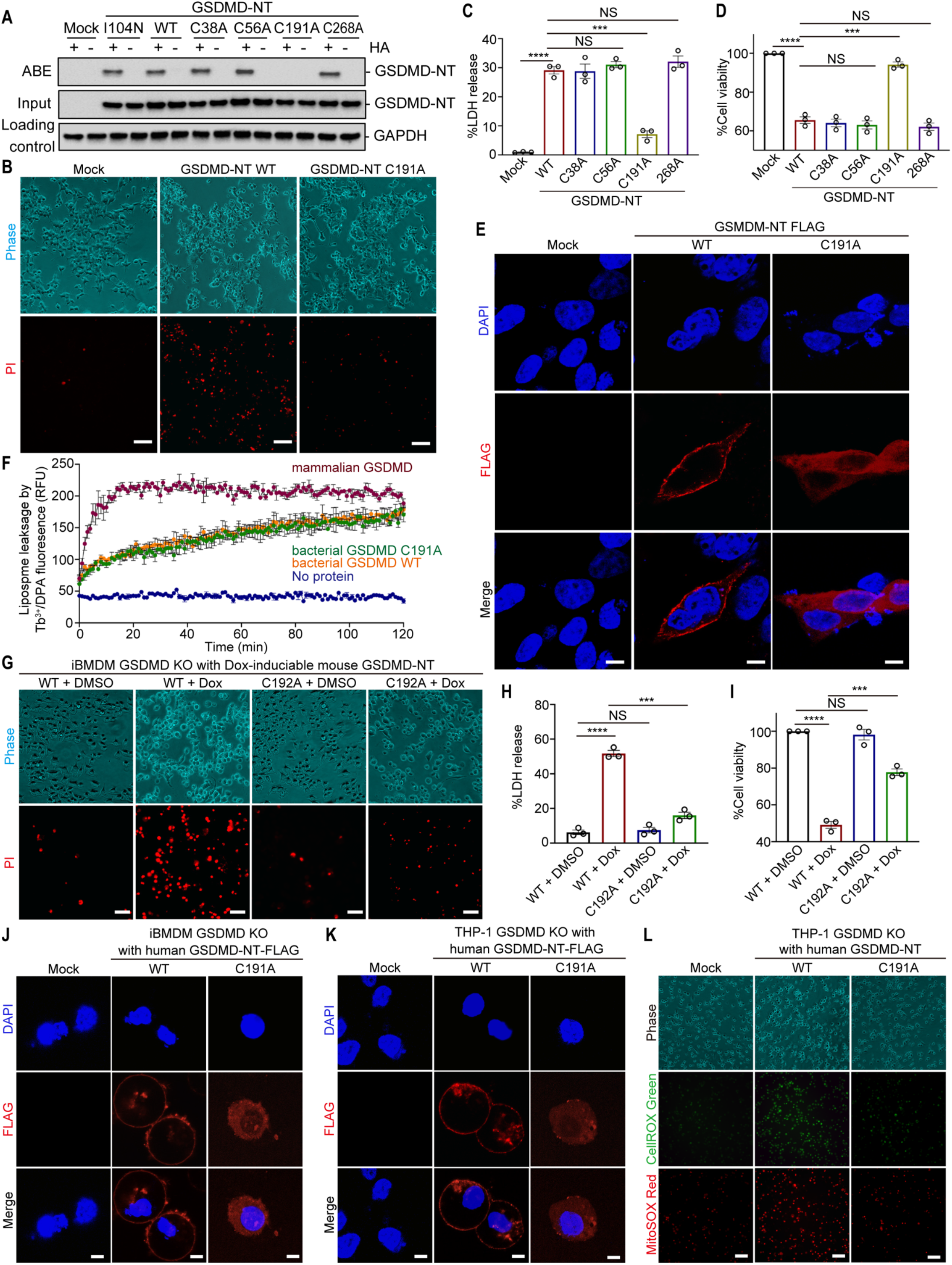
GSDMD is specifically palmitoylated at Cys191/192 (human/mouse) and defectiveness in palmitoylation compromises liposome leakage *in vitro*, and GSDMD membrane localization and pyroptosis in cells. (**A**) WT and mutant GSDMD-NT palmitoylation when overexpressed in HEK293T cells. Only the C191A mutant is defective in palmitoylation. (**B-D**). PI staining (B), LDH release (C), and viability (D) of HEK293T cells overexpressing WT or C191A GSDMD-NT, showing the impairment of C191A mutant in inducing pyroptosis. (**E**) Anti-Flag fluorescence imaging of HEK293T cells overexpressing WT or C191A GSDMD-NT, showing the cell membrane localization of WT GSDMD-NT and the diffuse cytoplasmic localization of C191A. Nuclei are marked by the DNA dye DAPI. (**F**) Markedly enhanced rate (∼10-fold) of liposome leakage by precleaved, Expi293 cell-expressed GSDMD in comparison to precleaved, bacterium-expressed WT GSDMD and the C191A mutant. (**G-I**) PI staining (G), LDH release (H), and viability (I) of GSDMD KO iBMDMs reconstituted with dox-inducible WT or C192A mouse GSDMD-NT, showing the impairment of C192A in inducing pyroptosis. (**J, K**) Anti-Flag fluorescence imaging of GSDMD KO iBMDMs (J) or GSDMD KO THP-1 cells (K) electroporated with WT or C191A human GSDMD-NT-FLAG, showing the cell membrane localization of WT GSDMD-NT and the diffuse cytoplasmic localization of C191A. (**L**) Measurement of cellular oxidative stress by CellROX and mitochondrial ROS by MitoSOX in GSDMD KO THP-1 cells electroporated with WT or C191A human GSDMD-NT. In (E), (J) and (K), nuclei are marked by the DNA dye DAPI. All results were obtained from at least 3 independent experiments. Scale bars represent 50 μm (B, G), 80 μm (L), and 5 μm (E, J, K). Error bars represent standard error of the mean (SEM). Statistics were measured by Student’s t-tests with *** for p<0.001, **** for p<0.0001, and NS (non-significant) for p>0.05.

The functional effects of palmitoylation revealed by the C191A mutation were quite striking, which prompted us to test if palmitoylated recombinant GSDMD also exhibits a higher ability to cause liposomes to release pre-encapsulated Tb^3+^ dye *in vitro*. We precleaved WT GSDMD expressed and purified from bacteria (non-palmitoylated) or Expi293 cells (partially palmitoylated), and C191A GSDMD from bacteria, and mixed them with liposomes to initiate pore formation. Strikingly, the Expi293 cell-expressed GSDMD induced faster liposome leakage than bacterium-expressed WT or C191A GSDMD at the same concentrations, ∼10-fold faster in the estimated initial rate at the first few minutes of this assay (**Fig. 2F**). This rate increase should be a lower limit since ABE on the same Expi293 cell-expressed GSDMD sample showed a percentage of palmitoylation at ∼27% (**fig. S2B**).

To investigate if the C191A mutant of GSDMD is also impaired in its function in macrophages, we used a previously established GSDMD KO immortalized mouse bone-marrow derived macrophage (iBMDM) line reconstituted with doxycycline (Dox)-inducible expression WT or C192A mouse GSDMD-NT (*23*). While WT GSDMD-NT reconstituted iBMDMs showed strong PI-positivity upon Dox induction, those reconstituted with C191A GSDMD-NT exhibited reduced PI positivity (**Fig. 2G, fig. S2C**). Similarly, LDH release was decreased (**Fig. 2H**), but cell viability by CellTiter-Glo was increased in the C191A GSDMD-NT iBMDMs (**Fig. 2I**). Immunofluorescence further showed that while WT GSDMD-NT reconstituted iBMDMs displayed strong cell membrane punctate staining indicative of active GSDMD, C191A mutant reconstituted iBMDMs showed more diffuse cytoplasmic staining (**Fig. 2J**).

We observed similar effects in GSDMD KO THP-1 cells electroporated with WT versus C191A GSDMD-NT, in which the WT GSDMD-NT was localized mainly on the cell membrane while the C191A mutant showed a largely diffuse and cytosolic distribution (**Fig. 2K**). Interestingly, the expression of WT, but not C191A GSDMD-NT, elevated cellular oxidative stress shown by CellROX, and mitochondrial reactive oxygen species (ROS) shown by MitoSOX (**Fig. 2L, fig. S2D, E**). These data may suggest that GSDMD pore formation itself increases mitochondrial and cellular ROS, the mechanism of which is unclear but may involve the damage of mitochondria or other organelles by GSDMD pores supported by the binding of GSDMD-NT to cardiolipin, a mitochondrial lipid (*3*). As shown below, cellular ROS conversely promotes GSDMD palmitoylation and pore formation, suggesting that these two processes are coupled and reinforce each other.

### GSDMD palmitoylation is regulated by reactive oxygen species

Unlike N-myristoylation, S-palmitoylation is mediated by a thioester bond between the modified Cys thiol group and the lipid chain, which makes it reversible by hydrolysis (at basic pH), attack from nucleophiles such as free thiols, or depalmitoylating enzymes. Because free thiol-containing reducing compounds are often used to remove a target thioester by transesterification (*30, 31*), and oxidative stress should decrease free thiols in cells, we hypothesized that GSDMD palmitoylation may be regulated by ROS or oxidative stress.

We selected the mitochondrial complex I inhibitor rotenone (Rot) and complex III inhibitor antimycin A (AMA), which have been shown to alter the electron transport chain in a way to drive ROS production (*21*), and N-acetylcysteine (NAC), a thiol-containing ROS quencher that modulates the intracellular redox state. We first transfected HEK293T cells with GSDMD-NT and treated the cells with these redox modulators, using untreated cells and 2-BP-treated cells as controls. In comparison with untreated cells, rotenone and antimycin A significantly increased and NAC significantly decreased GSDMD-NT palmitoylation (**Fig. 3A, B)**. PI positivity (**Fig. 3C, fig. S3A**) and LDH release (**Fig. 3D**) were increased in rotenone or antimycin A-treated cells but were reduced with NAC treatment. Cell viability on the other hand was decreased by rotenone or antimycin A treatment, but was enhanced by NAC treatment (**Fig. 3E**). Consistently, membrane localization of GSDMD-NT was increased in rotenone or antimycin A-treated cells and reduced in NAC-treated cells (**Fig. 3F**). In all experiments, 2-BP treatment decreased GSDMD-NT palmitoylation, cell death and membrane localization (**Fig. 3A-F**). By contrast, HEK293T-expressed ASC did not show differences in its cellular localization by treatment with 2-BP (**fig. S3B**).

**Fig. 3.**
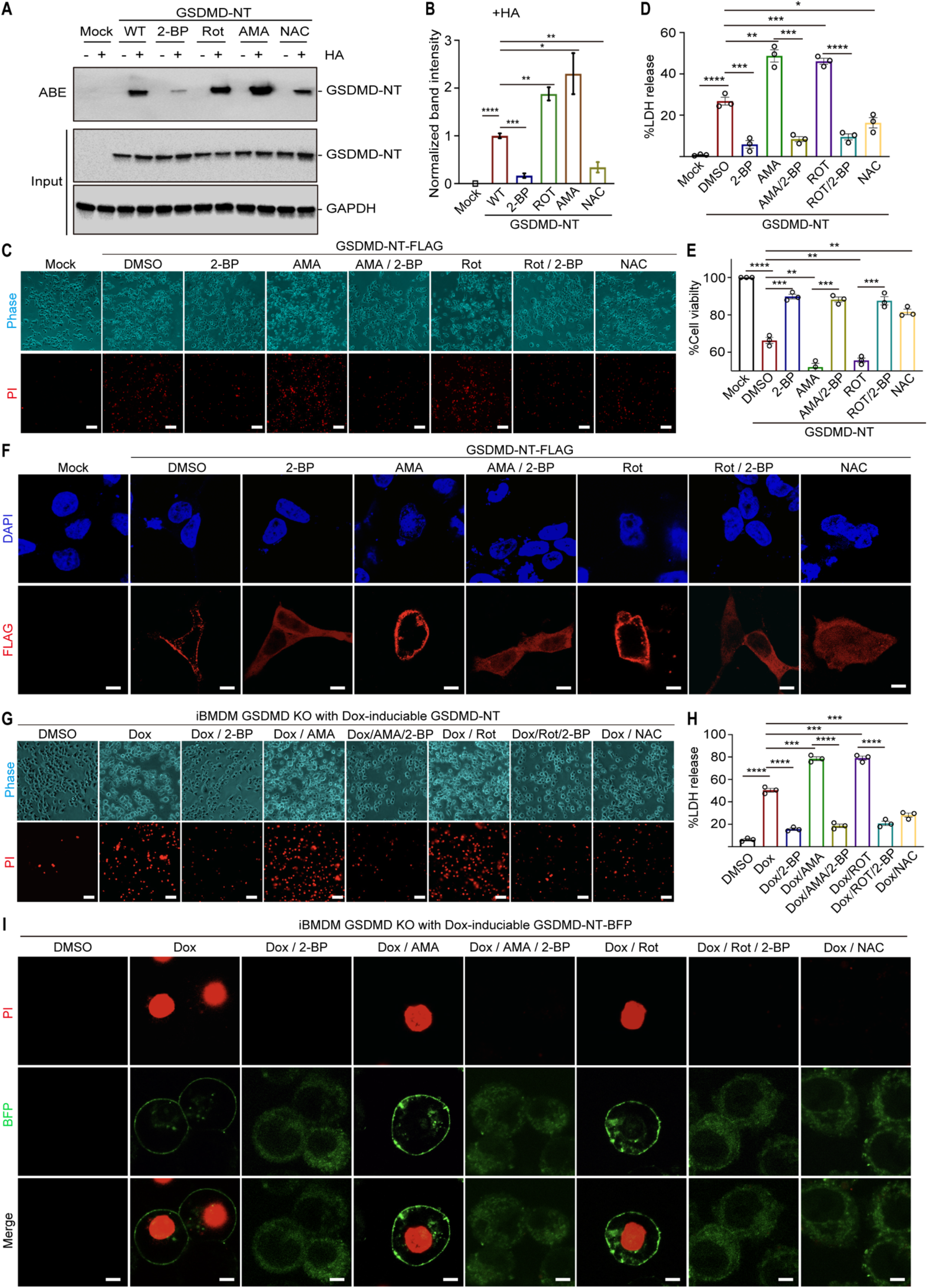
Regulation of GSDMD palmitoylation and pyroptosis by ROS modulators in HEK293T cells and iBMDMs. (**A, B**) GSDMD-NT palmitoylation detected by ABE when overexpressed in HEK293T cells and treated with ROS activators rotenone (Rot) and antimycin A (AMA) and ROS quencher N-acetyl cysteine (NAC) (A), showing the enhancement by Rot and AMA and suppression by NAC (B). (**C-E**). PI staining (C), LDH release (D), and viability (E) of HEK293T cells overexpressing GSDMD-NT and treated with ROS modulators showing enhanced pyroptosis by Rot and AMA, and decreased pyroptosis by NAC. (**F**) Anti-Flag fluorescence imaging of HEK293T cells overexpressing GSDMD-NT and treated with ROS modulators, showing increased cell membrane localization by Rot and AMA and its inhibition by 2-BP, as well as the diffuse cytoplasmic localization upon treatment by NAC. Nuclei are marked by the DNA dye DAPI. (**G, H**) PI staining (G) and LDH release (H) of GSDMD KO iBMDMs reconstituted with dox-inducible GSDMD-NT, showing enhanced pyroptosis by Rot and AMA and its inhibition by 2-BP, and decreased pyroptosis by NAC. (**I**) BFP fluorescence imaging of GSDMD KO iBMDMs reconstituted with dox-inducible GSDMD-NT-BFP and treated with ROS modulators, showing increased cell membrane localization by Rot and AMA and its inhibition by 2-BP, as well as the diffuse cytoplasmic localization upon treatment by NAC. PI staining is also shown to mark pyroptotic cells. All results were obtained from at least 3 independent experiments. Scale bars represent 50 μm (C, G), and 5 μm (F, I). Error bars represent SEM. Statistics were measured by Student’s t-tests with ** for p<0.01, *** for p<0.001, **** for p<0.0001, and NS (non-significant) for p>0.05.

We then used the GSDMD KO mouse iBMDM line reconstituted with Dox-inducible expression of WT GSDMD-NT. Rotenone or antimycin A treatment increased PI positivity (**Fig. 3G, fig. S3C, D**) and LDH release (**Fig. 3H**), but markedly decreased cell viability (**fig. S3E**) in Dox-induced cells. However, NAC treatment decreased PI positivity (**Fig. 3G, fig. S3C, D**) and LDH release (**Fig. 3H**), but markedly increased cell viability (**fig. S3E**) in Dox-induced cells. Punctate cell membrane localization of GSDMD-NT was increased in rotenone or antimycin A-treated cells and reduced in NAC-treated cells relative to Dox alone (**Fig. 3I**).

To further associate the regulation of palmitoylation with ROS, we performed ABE and measured ROS in THP-1 cells treated with LPS plus nigericin, pretreated or not with ROS generators or quenchers. In comparison with treatment by LPS plus nigericin alone, rotenone or antimycin A treatment significantly increased GSDMD palmitoylation, while NAC and the mTOR inhibitor Torin-1 which was shown previously to reduce ROS (*22*) decreased GSDMD palmitoylation **(Fig. 4A, B)**. ROS levels showed the same trend, with rotenone or antimycin A elevating, and NAC and the mitochondrial ROS scavenger MitoTEMPO (MitoT) reducing ROS (**Fig. 4C-E, fig. S4A**), supporting that manipulation of ROS could direct modulate GSDMD palmitoylation. Under these conditions, PI staining (**Fig. 4F, G, fig. S4B**) and LDH release (**Fig. 4H**) were significantly increased by rotenone or antimycin A and decreased by NAC, Torin-1, and MitoT, while cell viability had the inverse relationship (**Fig. 4I**). IL-1β release was also elevated by rotenone or antimycin A and decreased by NAC, Torin-1 and MitoT (**Fig. 4J**). Anti-GSDMD immunofluorescence imaging further confirmed increased GSDMD membrane localization by rotenone or antimycin A and largely diffuse GSDMD distribution by NAC (**fig. S4C**). While the effects of ROS augmenters were evident, the effect from a ROS quencher was almost on par with the 2-BP palmitoylation inhibitor, suggesting both the close coupling between ROS and palmitoylation and the requirement of palmitoylation for GSDMD pore formation and cell death induction.

**Fig. 4.**
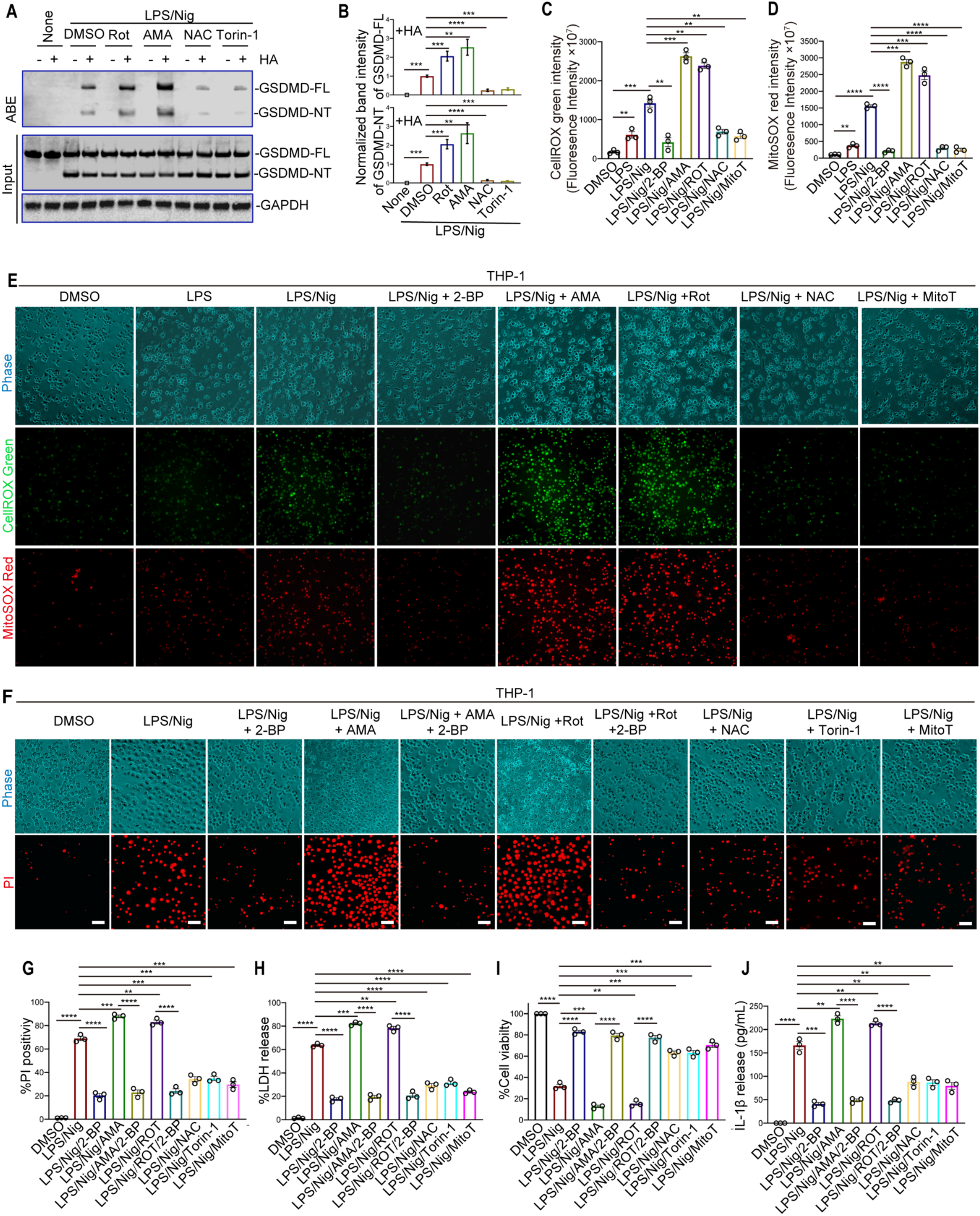
Regulation of GSDMD palmitoylation and pyroptosis by ROS modulators in THP-1 cells. (**A, B**) GSDMD palmitoylation detected by ABE in THP-1 cells treated or not with ROS modulators and upon stimulation by LPS and nigericin (A), showing that Rot and AMA enhanced, while NAC and Torin-1 decreased, GSDMD palmitoylation (B). (**C-E**) Measurement of cellular oxidative stress by CellROX and mitochondrial ROS by MitoSOX under similar conditions in (A) by imaging and quantification. Cellular and mitochondrial oxidative stress levels were similarly enhanced by Rot and AMA, decreased by NAC and MitoT, and correlated well with GSDMD palmitoylation. (**F-J**) PI staining (F, G), LDH release (H), viability (I), and IL-1β secretion (J) of THP-1 cells under similar conditions in (A), showing increased cell death and IL-1β secretion by Rot and AMA, and decreased cell death and IL-1β secretion by NAC, Torin-1 and MitoT. All results were obtained from at least 3 independent experiments. Scale bars represent 80 μm (E), and 50 μm (F). Error bars represent SEM. Statistics were measured by Student’s t-tests with ** for p<0.01, *** for p<0.001, and **** for p<0.0001.

### GSDMD is palmitoylated primarily by zDHHC5 and zDHHC9

The thioesterification of palmitate to Cys residues on target proteins is catalyzed by Asp-His-His-Cys (zDHHC)-family palmitoyl S-acyltransferases with 23 members in humans (zDHHC1-9 and 11-24) (*32*). To elucidate which zDHHCs palmitoylate GSDMD, we co-expressed all 23 zDHHCs with GSDMD in HEK293T cells and performed click chemistry detected by rhodamine fluorescence (**Fig. 1B**). Probably because GSDMD is already significantly palmitoylated when transfected alone, co-expression with zDHHCs had relatively modest effects on GSDMD palmitoylation. Nonetheless, zDHHC1, 5, 9, 12, 17, 19-21 appeared to increase GSDMD palmitoylation (**Fig. 5A, fig. S5A**). We then checked the expressions of these zDHHCs in THP-1 and HEK293T cells using the Human Protein Atlas database and found that only zDHHC5, 9, 12, 17, and 20 are significantly expressed (**Fig. 5B**).

**Fig. 5.**
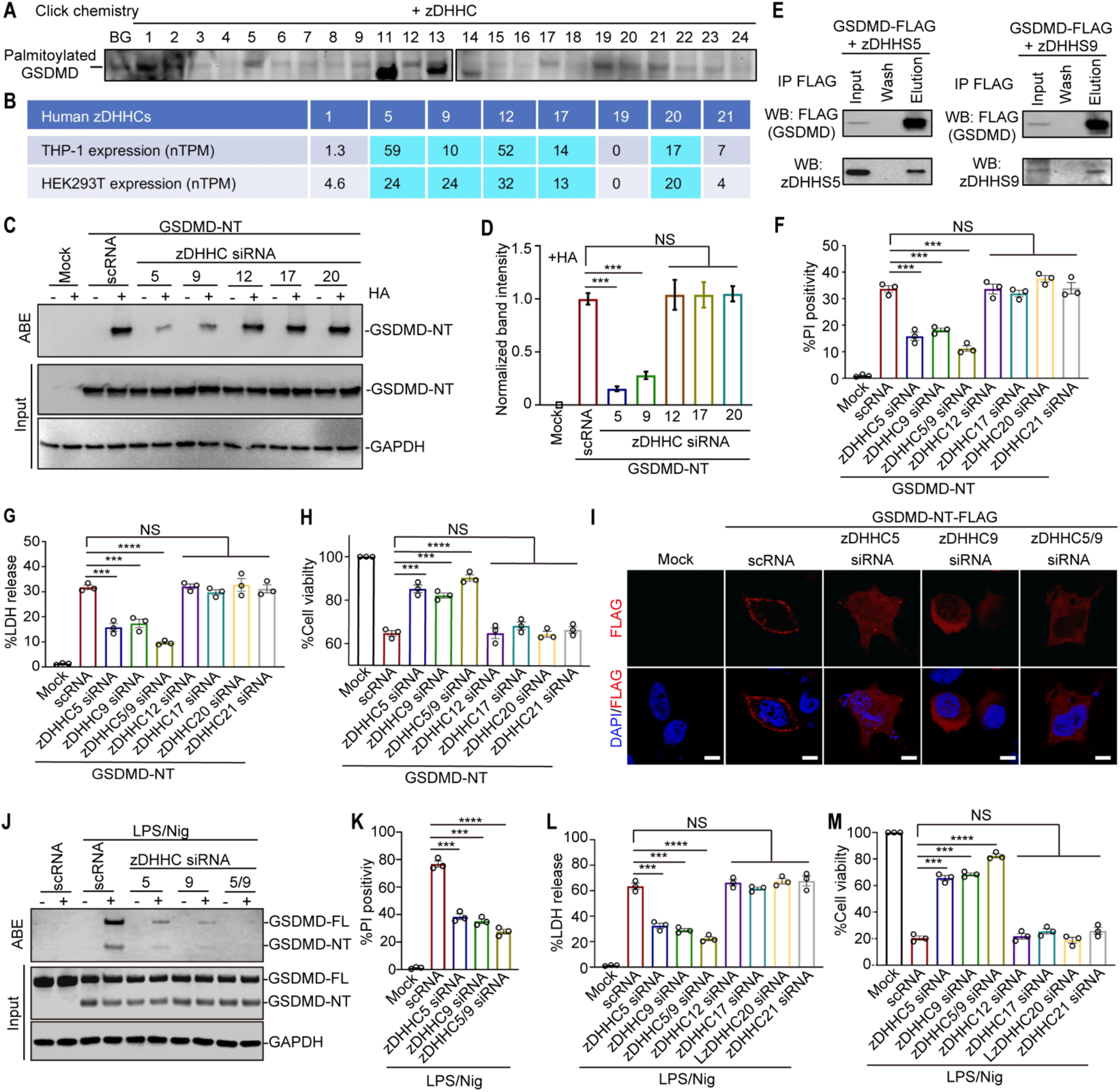
Identification of zDHHC5 and zDHHC9 as the main palmitoyltransferases for GSDMD palmitoylation. (**A**) Click chemistry screen by rhodamine labeling (**Fig. 1B**) in 293T cells co-expressing the 23 human zDHHCs with GSDMD to identify zDHHCs that enhanced GSDMD palmitoylation. Additional bands in the blots are likely from autopalmitoylation of zDHHCs. (**B**) Expression data of zDHHCs in HEK239T and THP-1 cells from Protein Atlas, identifying 5 highly expressed zDHHCs among those that enhanced GSDMD palmitoylation. (**C, D**) GSDMD-NT palmitoylation detected by ABE in HEK293T cells upon siRNA knockdown of the 5 highly expressed zDHHCs or a scrambled siRNA (scRNA) as a control (C), showing that knockdown of zDHHC5 and zDHHC9 compromised GSDMD-NT palmitoylation (D). (**E**) Anti-FLAG pulldown of zDHHS5 (left, catalytic mutant of zDHHC5) and zDHHC9 (right) by co-expressed GSDMD-FLAG. (**F-H**) PI positivity (F), LDH release (G), and viability (H) of GSDMD-NT expressing HEK293T cells upon siRNA knockdowns of zDHHCs. (**I**) Anti-FLAG immunofluorescence imaging of HEK293T cells with siRNA knockdowns of zDHHC5, zDHHC9, or both overexpressing GSDMD-NT-FLAG. Only cells treated with scRNA showed strong cell surface staining. Nuclei are marked by the DNA dye DAPI. (**J**) GSDMD palmitoylation detected by ABE in THP-1 cells upon siRNA knockdown of zDHHC5, zDHHC9, or both and treatment with LPS plus nigericin, showing that these knockdowns compromised GSDMD palmitoylation. (**K-M**) PI positivity (K), LDH release (L), and viability (M) of THP-1 cells upon siRNA knockdown of zDHHC5, zDHHC9, both, or other zDHHCs and treatment with LPS plus nigericin. All results were obtained from at least 3 independent experiments. Scale bars represent 5 μm (I). Error bars represent SEM. Statistics were measured by Student’s t-tests with *** for p<0.001, **** for p<0.0001, and NS (non-significant) for p>0.05.

We thus used siRNAs to knock down these 5 zDHHCs in HEK293T cells when transfecting GSDMD-NT. We found that zDHHC5 and to a lesser extent zDHHC9 were responsible for GSDMD-NT palmitoylation; siRNAs to other zDHHCs or scrambled siRNA (scRNA) did not have observable effects (**Fig. 5C, D**). Most human zDHHCs have been shown to localize to the early biosynthetic pathway (endoplasmic reticulum and Golgi), but zDHHC5 appears primarily localized to the plasma membrane (*33-36*). Our co-expression of zDHHC5 and zDHHC9 with GSDMD confirmed these potential localizations (**fig. S5B**). We further validated the interaction of GSDMD-FLAG with zDHHC5 and zDHHC9 by anti-FLAG co-immunoprecipitation (IP), and zDHHC5 and zDHHC9 were not pulled down without co-expression with the GSDMD-FLAG bait (**Fig. 5E, fig. S5C**). The same siRNA knockdown of zDHHC5 or zDHHC9, and more significantly the double knockdown of zDHHC5 and zDHHC9, but not the knockdown by scRNA or other zDHHCs, decreased PI positivity and LDH release (**Fig. 5F, G**), and increased cell viability (**Fig. 5H**). Cellular imaging of GSDMD-NT also revealed the impairment of membrane localization upon siRNA knockdown of zDHHC5, zDHHC9, or both (**Fig. 5I**).

Similarly, knockdown of zDHHC5, zDHHC9, or both in THP-1 cells followed by LPS plus nigericin stimulation decreased palmitoylation without affecting GSDMD cleavage (**Fig. 5J, fig. S6A**), reduced PI positivity and LDH release (**Fig. 5K, L**) and increased cell viability (**Fig. 5M**). Anti-GSDMD immunofluorescence displayed its cell membrane localization in scRNA-treated THP-1 cells, but more diffuse and cytosolic staining upon treatment by siRNA of zDHHC5, zDHHC9, or both (**fig. S6B**).

### Mechanism for ROS-dependent GSDMD palmitoylation

We wondered how GSDMD palmitoylation is regulated by ROS. Given the strong inhibitory effects on GSDMD palmitoylation by 2-BP under all conditions tested, we first examined if palmitoyltransferases are upregulated by inflammasome activation which is associated with increased ROS, or by direct treatment with ROS modulators. We found by immunoblot that the level of zDHHC5, the main palmitoyltransferase we identified, remained essentially constant in THP-1 cells upon treatment by LPS, LPS plus nigericin, or with added ROS activators or NAC (**Fig. 6A**). This experiment suggested that the enhanced production of palmitoylated GSDMD by zDHHC5 and other palmitoyltransferases is not due to upregulation of zDHHC expression, but either enhanced substrate availability (e.g. palmitoyl-CoA) or enzymatic activity.

**Fig. 6.**
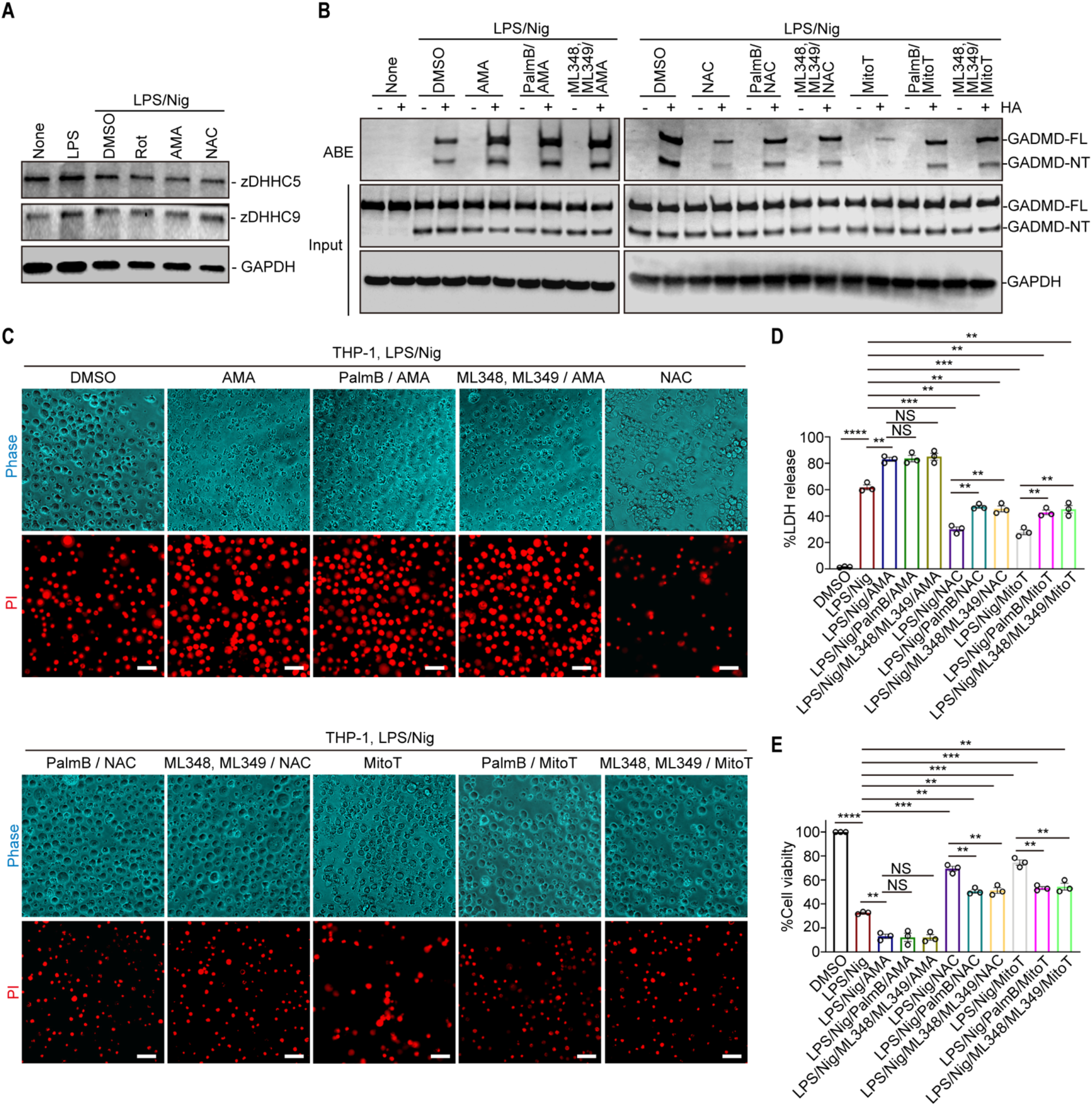
Mechanism of ROS-regulated GSDMD palmitoylation. (**A**) Expression level of zDHHC5 and zDHHC9 in THP-1 cells treated with LPS, LPS plus nigericin, or LPS plus nigericin additionally treated with ROS modulators. No apparent changes were observed. (**B**) GSDMD palmitoylation detected by ABE in THP-1 cells pre-treated by PalmB (general inhibitor), or ML348 (APT1 inhibitor)/ML349 (APT2 inhibitor) together, or not, followed by addition of an ROS modulator, LPS and nigericin. PalmB or ML348/ML349 did not alter GSDMD palmitoylation under treatment of the ROS activator AMA, but significantly increased GSDMD palmitoylation under treatment of ROS quenchers NAC and MitoT. (**C-E**) PI staining (C), LDH release (D) and cell viability (E) of the same treatment as in (B). Scale bars in (C) represent 30 μm. All results were obtained from at least 3 independent experiments. Error bars represent SEM. Statistics were measured by Student’s t-tests with ** for p<0.01, *** for p<0.001, **** for p<0.0001, and NS (non-significant) for p>0.05.

To address if the changes in palmitoylation under different conditions are influenced by changes in the action of depalmitoylating enzymes (*37*), we used inhibitors for these enzymes. While palmostatin B (PalmB, K_i_ = 0.67 µM) is a promiscuous depalmitoylase inhibitor, ML348 and ML349 are more potent isoform-selective inhibitors of acyl protein thioesterase 1 (APT1, also known as LYPLA1, K_i_ = 280 nM) and APT2 (also known as LYPLA2, K_i_ = 120 nM), respectively (*37-40*). Pre-treatment by PalmB or ML348/ML349 together, or not, followed by addition of the ROS activator antimycin A, LPS and nigericin did not alter GSDMD palmitoylation (**Fig. 6B**), PI positivity (**Fig. 6C, Fig. S7A, B**), LDH release (**Fig. 6D**) or cell viability (**Fig. 6E**). By contrast, the same treatment regimen of depalmitoylating enzyme inhibitors, but with ROS activator replaced by the ROS quencher NAC or the mitochondrial ROS scavenger MitoTEMPO (MitoT), significantly but moderately increased GSDMD palmitoylation, PI positivity and LDH release (**Fig. 6B-D**), and decreased cell viability (**Fig. 6E**), but did not restore cell death to the level without any ROS modulators (**Fig. 6B-E**). These data support that depalmitoylation may have already been inhibited under high ROS to contribute to increased GSDMD palmitoylation.

## Discussion

Our studies demonstrate that GSDMD palmitoylation, a form of lipidation, is an obligate requirement for its membrane pore formation in addition to proteolytic cleavage, which may also serve as a general principle for activation of the GSDM family suggested by ubiquitous palmitoylation of GSDM-NTs. Because palmitoylation is the only reversible lipid modification, its use provides a regulated checkpoint for the potentially devasting effect of GSDMD-mediated pyroptosis. We propose the following model for how palmitoylation enhances GSDMD-NT pore formation by enabling membrane insertion and also accelerating membrane localization and oligomerization (**Fig. 7**). First, GSDMD is not significantly palmitoylated in a resting state due to low ROS. Upon inflammasome activation, ROS level is massively upregulated with concomitant surge in palmitoylation of both GSDMD-FL and cleaved GSDMD. Other cellular stress could further enhance ROS and the level of GSDMD palmitoylation. Palmitoylated GSDMD-FL remains cytosolic in cells, likely by folding back its palmitoyl group to the protein to make it unavailable for membrane interaction, whereas cleaved, palmitoylated GSDMD quickly localizes to the cell membrane, displaces the CT while the NT oligomerizes and inserts to form transmembrane pores. Cleaved GSDMD that is yet to be palmitoylated should have its membrane-binding patch exposed and thus in theory could localize to the membrane, displace its CT and oligomerize into GSDMD-NT prepores (*8*). This hypothesis is consistent with the previous observation that sometimes GSDMD-NT can localize as oligomers to the cell membrane without causing cell death (*22, 41*). We posit that membrane-bound unpalmitoylated GSDMD-NT is palmitoylated by cell surface localized zDHHC5, which triggers GSDMD-NT prepore to pore transition, and pyroptosis. Furthermore, although we do not have experimental evidence, the ring of membrane-inserted lipid acyl chains at the membrane-facing side of the GSDMD-NT pore may cause phase separation of surrounding lipids, like the effect by S-palmitoylated SARS-CoV-2 spike protein (*42*), and perhaps induce GSDMD-NT pore clustering that is suggested by the punctate features in GSDMD-NT membrane localization. Overexpression of GSDMD-NT, which increases ROS, could similarly cause pore formation and pyroptosis.

**Fig. 7.**
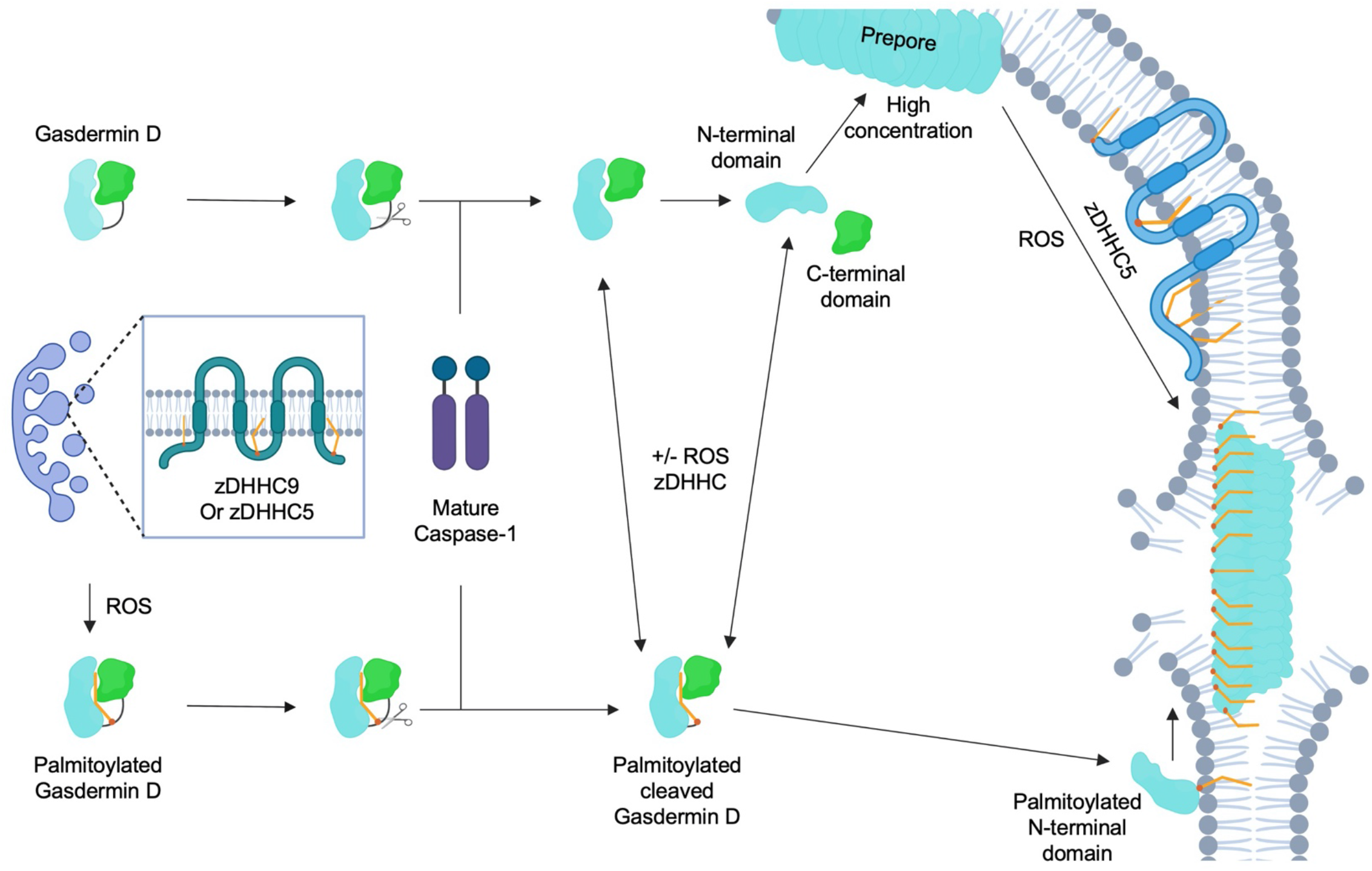
A proposed model for how palmitoylation enhances GSDMD-NT pore formation. See text for details.

Our studies reveal a specific effect on palmitoylation downstream of the rather non-specific ROS, which makes biological sense. They affirm the critical point that oxidative stress directly promotes GSDMD palmitoylation, independent of the upstream inflammasome pathway and GSDMD cleavage, and explains the key role of ROS in GSDND-NT pore formation (*22, 23*). That palmitoylation appears to be a common modification of the GSDM family may further implicate that other GSDM-mediated pyroptosis is also dependent on cellular oxidative stress, making pyroptosis a crucial downstream cellular consequence to high ROS. We do not yet know the exact mechanism by which ROS enhances the level of GSDMD palmitoylation, but high ROS appears to both increase the palmitoylation reaction shown by 2-BP effect, and decrease the depalmitoylation reaction shown by lack of effect from depalmitoylase inhibitors. We speculate, but have no evidence, that high ROS may increase substrate flux to zDHHCs to increase the production of palmitoylated GSDMD. In addition, high ROS may reduce non-enzymatic depalmitoylation by, among other things, a reduction in free thiols in cells that can break thioester bonds by transesterification (*30, 31*).

The Cys191/192 residue is also the target for other modifications including oxidation (*23*), and the target of multiple small molecules that inhibit GSDMD functions, including necrosulfonamide (NSA), disulfiram (DSF) and dimethyl fumarate (DMF) (*43-45*). It is templating to speculate that the mechanism of inhibition by these small molecules in cells may be related to competition with GSDMD palmitoylation at the Cys191/192 site. Endogenous fumarate that causes GSDMD succination may even be an intrinsic regulation to depalmitoylate GSDMD and thus deactivate GSDMD-mediated pore formation (*45*). How the different modifications interplay in cells remains to be addressed and may reveal interesting new biology.

## Materials and Methods

### Constructs

Full-length human GSDMD containing an internal human rhinovirus 3C protease (3C) site was cloned into the pDB.His.MBP vector after the N-terminal His6-maltose binding protein (MBP) tag as previously described (GSDMD-MBP-3C) (*8*) and the C191A mutant was generated. GSDMD-FL, GSDMD-NT, and GSDMD-CT were amplified by polymerase chain reaction and subcloned into a pLV vector containing C-terminal mCherry (GSDMD-mCherry) or the pcDNA3.1 vector tagged with C-terminal FLAG (GSDMD-FLAG). GSDMA-FL, GSDMB-FL, GSDMC-FL, GSDME-FL, and GSDME-NT were cloned into the pCMV4 vector with N-terminal FLAG. All plasmids were transformed into DH5α-competent cells, subsequently mini-prepped (Qiagen), and verified by sequencing. HeaTil screen (human zDHHC screen) plasmids pLH1-24 have been described previously (*32*). The caspase-1 construct that co-expresses p20 and p10 subunits was a gift from T. Sam Xiao. All mutations in this study were introduced using the QuikChange II XL site-directed mutagenesis kit (Agilent Technologies, Cat #200521), KLD Enzyme Mix, or Gibson Assembly Master Mix (New England Biolabs).

### Cell lines

Human monocytic THP-1 and Human embryonic kidney 293T cells (HEK293T) cell lines were purchased from ATCC. GSDMD KO THP-1 cells are a gift from Daniel Bachovchin. GSDMD KO immortalized mouse bone marrow–derived macrophages (iBMDMs) reconstituted with Dox-inducible GSDMD-NT WT and the C192A mutant are as described (*22, 23*).

### Mammalian cell culture and transfection

HEK293T and iBMDMs were cultured in Dulbecco’s modified Eagle’s medium with glutamine (DMEM, GIBCO, ThermoFisher Scientific) supplemented with 10% fetal bovine serum (FBS, Sigma, TMS-013-B), 1% penicillin/streptomycin mix (GIBCO, ThermoFisher Scientific), L-glutamine (GIBCO, ThermoFisher Scientific) and sodium pyruvate (GIBCO, ThermoFisher Scientific). THP-1 cells were maintained in Roswell Park Memorial Institute medium (RPMI1640, GIBICO, ThermoFisher Scientific) with L-glutamine, supplemented with 10% FBS and 1% penicillin/streptomycin mix (GIBCO, ThermoFisher Scientific). They were differentiated by treatment with 200 ng/mL phorbol 12-myristate 13-acetate (PMA, Sigma-Aldrich, P8139–5MG) for 48 hours. All cells were propagated in a humidified incubator at 37 °C and with 5% CO_2_.

### Expression and purification of GSDMD proteins in bacteria and Expi293 cells

GSDMD-MBP-3C WT and C191A mutant constructs were transformed into *E. coli* BL21 (DE3) cells, plated and incubated overnight at 37 °C. Single colonies were picked the next day and grown in Luria Broth (LB) media supplemented with 50 μg/mL kanamycin at 37 °C. The cultures (Agilent Technologies, 230280) were induced at an optical density at 600 nm (OD600) of 0.6 by 1 mM isopropyl β-d-1-thiogalactopyranoside (IPTG) and incubated for 18 hours at 18 °C before collection. Cells were pelleted by centrifugation at 4,000 g for 30 min and resuspended in buffer A (40 mM HEPES at pH 7.0, 150 mM NaCl) supplemented with 5 mM imidazole for lysis by sonication. His6-MBP-tagged GSDMD was enriched on Ni-NTA beads and eluted by buffer A supplemented with 500 mM imidazole. The His6-MBP tag was cleaved by His6-tagged tobacco etch virus (TEV) protease at 4 °C overnight. His6-MBP and His6-TEV were removed using a Ni-NTA column and the flow-through containing GSDMD was further purified using a Superdex 200 Increase 10/300 GL (Cytiva) size-exclusion column equilibrated with buffer A. Peak fractions were collected, analyzed via SDS-PAGE for purity, and snap frozen for liposome leakage assays.

The GSDMD-FLAG construct was transfected to Expi293F cells that were maintained in 1000 ml Expi293 Expression Media (Thermo Fisher), fed with 6 mM KCl and grown to 2.5 × 10^6^ cells/ml, using polyethylenimine (PEI, Polysciences, Inc.). The cells were fed with 10 mM sodium butyrate and 10 ml 45% D-(+) glucose solution 12 hours after transfection. The cells were grown for another 2 days and harvested by centrifugation at 4,000 g for 30 min. The cell pellet was resuspended in buffer A and lysed by sonication (2-s on, 8-s off, 3.5 min total on, 40% power), and centrifuged at 40,000 rpm for 1 hour. The supernatant was collected and incubated with ANTI-FLAG® M2 Affinity Gel (Sigma-Aldrich, A2220-4X25ML) for overnight at 4 °C with gentle rotation. After washing, the protein was eluted using buffer A with 150 ng/µL 3X FLAG peptide (sigma, F4799-25MG). The eluted GSDMD protein was snap frozen for other assays.

### Acyl Biotin Exchange (ABE)

Cells were treated and collected as specified and spun down. Cell pellets were resuspended in 0.2 mL buffer A (50 mM Tris-HCl pH 7.4, 1 mM EDTA, 150 mM NaCl, 1% NP-40) with 20 mM N-ethylmaleimide (NEM, Sigma, E3876-5G) and protease inhibitor cocktail. Samples were incubated for 2 hours with gentle rotation at 4 °C and centrifuged at 20,000 g for 20 min. The supernatant was collected and Pierce™ BCA Protein Assay Kit (ThermoFisher, 23225) was used to determine the protein concentration to equalize the amount of protein in each sample.

The sample was then precipitated with a methanol: chloroform: water mixture. Briefly, 800 µL methanol, 300 µL chloroform, and 600 µL ddH_2_O were added to each sample sequentially with thorough mixing by vortex at each step. The mixtures were spun down for 5 minutes at 16,000 g in a microfuge and the top aqueous layer was discarded. 800 µL methanol was added to resuspend each pellet. The mixtures were thoroughly vortexed and spun down for 5 minutes at 16,000 g, and the supernatants were removed as much as possible. The pellets were air-dried for 15-20 minutes. The above methanol/chloroform precipitation was repeated two additional times.

The dried pellets were resuspended in 0.2 mL buffer B (50 mM Tris-HCl pH 7.4, 5 mM EDTA, 150 mM NaCl, 1% NP-40, 1% SDS) with or without 0.7 M hydroxylamine (50% by weight in H_2_O, Sigma, 438227-50ML) to totally dissolve the pellets by pipetting or even water sonication. 20 µL 10 mM N-[6-(biotinamido)hexyl]-3′-(2′-pyridyl dithio)propionamide (biotin-HPDP, Cayman Chemical Company, 16459) in DMSO was added to each sample, mixed gently and incubated with gentle rotation at room temperature for 1 hour. The samples were precipitated by the same above methanol/chloroform protocol and dissolved in buffer B with 1 mM biotin-HPDP and incubated with gentle rotation at room temperature. After incubation for 1 hour, the samples were precipitated by the same above methanol: chloroform protocol three times.

After drying, the pellets were resuspended in 0.1 mL buffer B, and 20 µL of each sample was taken out as a loading control and combined with 20 µL 2 x SDA-PAGE Laemmli buffer. The SDS concentration in the remaining samples were diluted 10-fold to 0.1% by buffer C (50 mM Tris-HCl pH 7.4, 1 mM EDTA, 150 mM NaCl) before adding 30 µL prewashed streptavidin agarose. The mixture was incubated overnight at 4 °C with rotation. The agarose beads were washed three times using buffer C with 0.1% NP-40 and 0.1% SDS, mixed with 2x SDS-Laemmli buffer containing β-mercaptoethanol and incubated at 98 °C for 5 minutes before running an SDS-PAGE. The gels were then transferred and immunoblotted with appropriate antibodies.

### Cell viability and microscopy-based cytotoxicity assays

Lactate dehydrogenase (LDH) release was measured using LDH-Glo Cytotoxicity Assay kit (Promega, J2380) according to the manufacturer’s instructions. Cell viability was assessed by measuring ATP levels using the CellTiter-Glo Luminescent Cell Viability Assay (Promega, G7570) according to the manufacturer’s instructions. Luminescence was measured on a SYNERGY microplate reader (Biotek). For microscopy-based cytotoxicity analysis, cells were seeded in 24-well plates or CELLview Cell Culture Dish with four compartments) (USA Scientific, 5662-7870) in culture media with 1 μg/ml propidium iodide (PI, BD Bioscience, 556463) followed by calculating PI positivity on either an Inverted Nikon Ti2 fluorescence microscope with a 40X objective (N.A. = 1.2) with a stage top incubator to maintain 37 °C and 5% CO_2_ or on a Leica TCS SP8 Laser Scanning Confocal (Leica) fluorescence microscope with an incubator to maintain 37 °C and 5% CO_2_.

### Detection of human IL-1β by ELISA

Sandwich enzyme-linked immunosorbent assay detection (ELISA) kit for human IL-1β (Invitrogen, catalog # 88-7261-88) was used at the specified temperature and conditions according to the manufacturer’s instructions.

### siRNA assay

For HEK293T cells, scrambled siRNA (scRNA, control), or siRNA for zDHHCs were transfected into cells for 48 hours using Lipofectamine 3000 (Invitrogen, L3000-015) according to manufacturer’s instructions. For THP-1 cells, siRNAs were electroporated using the Neon™ Transfection System (Invitrogen) and the cells were grown for 48 hours before conducting any further experiments.

### Immunoblot analysis

Cells were collected and lysed using Lysis buffer (50 mM Tris-HCl pH 7.4, 150 mM NaCl, 1% NP40 supplemented with Halt protease inhibitor cocktail (Sigma, S8830-20Tab). Samples were incubated for 2 hours with gentle rotation at 4 °C and then centrifuged at 20,000 g for 20 min. The supernatant was collected and Pierce™ BCA Protein Assay Kit (Thermo, 23225) was used to determine the protein concentration to equalize the amounts of proteins in all samples. Samples were subjected to SDS-PAGE on 4-12% Tris-Glycine Gels (Bio Rad) and transferred to a 0.2 μm nitrocellulose membrane using the iBlot system (Invitrogen) followed by blocking with 5% Milk in Tris-buffered saline with Tween 20 (TBST) for 30 minutes. After blocking, the membrane was washed using TBST three times, 5 minutes each. For anti-FLAG immunoblot, the membrane was incubated with mouse monoclonal anti-FLAG® M2-peroxidase (HRP) antibody (Sigma, A8592) for 3 hours at room temperature or overnight at 4 °C. For other immunoblots, the membrane was incubated with a primary antibody at 4 °C overnight followed by extensive washing and incubation with a secondary antibody, 1:1000 HRP-goat anti-rabbit IgG (Invitrogen, 31460) or 1:1000 HRP-goat anti-mouse IgG (Invitrogen, 31432) for 3 hours at room temperature.

Immunoblots were probed with the following primary antibodies: 1:1000 rabbit anti-GSDMD monoclonal antibody (Cell Signaling Technology, 39754), 1:1000 rabbit anti-GSDMD polyclonal antibody (Novus Biologicals, NBP-33422), 1:500 rabbit anti-zDHHC5 polyclonal antibody (Proteintech, 21324-1-AP), 1:500 rabbit anti-zDHHC9 polyclonal Antibody (ABclonal, A7977), 1:1000 rabbit anti-calnexin Antibody (Cell Signaling Technology, 2433S), or 1:5000 mouse anti-GAPDH monoclonal antibody (Proteintech, 60004-1-IG). Visualization used SuperSignal™ West Atto Ultimate Sensitivity Substrate kit (Sigma, A38556). Western blot images were captured using a BioRad Chemidoc™ MP imaging system and band intensities were quantified using ImageJ.

### Immunofluorescence assays

HEK293T cells were plated in 24-well plates with poly-L-Lysine coated glass coverslips (12 mm) at 5 × 10^4^ cells per well. After 24 hours, these cells were pre-treated or not with 2-bromopalmitate (2-BP, Sigma, 21604-1G) at 50 μM for 30 min, followed by transfection of appropriate constructs using Lipofectamine 3000 (Invitrogen, Cat # L3000008). At 16 hours post-transfection, cells were treated or not with antimycin A (AMA, Sigma, A8674) at 10 μg/mL, Rotenone (ROT, Sigma, R8875-1G) at 10 μM, or N-acetyl cysteine (NAC, Sigma, A9165-25G) at 15 mM for 4 hours and fixed by addition of 500 µL 4% paraformaldehyde (PFA) in phosphate-buffered saline (PBS) at 37 °C for 15 min. After washing twice with PBS, cells were blocked with 2% bovine serum albumin (BSA) in PBS for 30 minutes at room temperature. Cells were then incubated with the L5 clone of Alexa Fluor® 594 anti-Flag antibody (BIOLEGEND INC, 637314), diluted 1:200 in PBS buffer containing 2% BSA, at 4 °C in the dark overnight. Following 3 washes with PBS, the coverslips were mounted with ProLong Gold Antifade Mountant with DAPI (Thermo Fisher Scientific, P36941) onto slides and left to dry for 1.5 hours in the dark.

For THP-1 GSDMD KO cells or GSDMD KO iBMDMs, cells were pre-treated or not with 2-BP (50 μM) for 30 min, followed by electroporation of appropriate constructs using the Neon™ Transfection System (Invitrogen). After electroporation, these cells were plated in 24-well plates with poly-L-Lysine coated glass coverslips (12 mm) at 5 × 10^4^ cells per well. At 16 hours post-electroporation, cells were treated or not with AMA (10 μg/mL), ROT (10 μM), NAC (15 mM), Torin-1 (Cayman Chemical Company, 10997) at 2 μM, or MitoTempo (MitoT, Sigma, SML0737) at 500 μM) for 4 hours and fixed, blocked and incubated with appropriate antibodies as described above for HEK293T cells.

For THP-1 WT cells, cells were plated in 24-well plates with poly-L-Lysine coated glass coverslips (12 mm) at 2.5 × 10^4^ cells per well and treated with 200 ng/mL PMA for 48 hours. Cells were pre-treated or not with 2-BP (50 μM) for 30 min, and treated with AMA (10 μg/mL), ROT (10 μM), NAC (15 mM), Torin-1 (2 μM), or MitoT (500 μM) for 4 hours, LPS (1 μg/mL) for 3 hours and nigericin (20 μM) for 1 hour. The cells were then fixed, blocked, and incubated with anti-GSDMD primary antibody at 4 °C overnight. After washing, cells were incubated with Alexa Fluor™ 568 goat anti-rabbit IgG (H+L) cross-adsorbed secondary antibody (Invitrogen, A11036) for 3 hours at room temperature. Following 3 washes with PBS, coverslips were mounted with ProLong Gold Antifade Mountant with DAPI (Thermo Fisher Scientific, Cat# P36941) onto slides and left to dry for 1.5 hours in the dark.

For GSDMD KO iBMDMs reconstituted with Dox-inducible, blue fluorescent protein (BFP)-tagged GSDMD-NT, cells were plated in CELLview Cell Culture Dish with four compartments at 5 × 10^4^ cells per compartment. After 24 hours, cells were treated or not with 2-BP for 30 minutes, Dox for 6 hours, and AMA (10 μg/mL), ROT (10 μM), NAC (15 mM), Torin-1 (2 μM), or MitoT (500 μM) for 4 hours. After treatment with 1 μg/mL PI for 30 minutes, imaging was performed.

All images were taken using a Leica TCS SP8 confocal laser scanning microscope at the Boston Children’s Hospital Microscopy Facility. The images were identically acquired and processed using Adobe Illustrator or ImageJ software.

### Live cell ROS measurements

Cells were manipulated and treated identically for immunofluorescence of WT THP-1 cells to activate the NLRP3 inflammasome under different conditions. 5 μM CellROX™ Green (Invitrogen, C10444) and 1 μM MitoSOX™ Red (Invitrogen, M36008) were added to the cells for 30 min at 37 °C. Images were taken on an Inverted Nikon Ti2 fluorescence microscope with a 40X objective (N.A. = 1.2) with a stage top incubator to maintain 37 °C and 5% CO_2_. The fluorescence intensities of CellROX™ Green and MitoSOX™ Red were determined by ImageJ software.

### Metabolic labeling *in cellulo*

Expression of GSDMD-mCherry was carried out by transfecting HEK293T cells with 2 μg DNA and Lipofectamine 2000. After 16 hours the media were replaced with 1 ml of DMEM conditioned with 1% charcoal-stripped fatty acid-free bovine serum albumin (BSA). The same media contained 50 μM 2-BP solubilized in DMSO for 2-BP experiments. Two hours later the cells were fed with 100 µM saponified fatty acid alkyne 17-Octadecynoic Acid (17-ODYA) or 25 µM 13-tetradecynoic acid (13-TDYA) dissolved in DMEM conditioned with 20% charcoal-stripped fatty acid-free BSA. After 6 hours cells were harvested into microfuge tubes with two washes of 1×PBS.

Cells were resuspended in radioimmunoprecipitation assay (RIPA) buffer supplemented with 100× protease inhibitor cocktail and benzonase nuclease, extracted for 90 minutes with rotation, and centrifuged at the max speed of a microfuge for 10 min at 4 °C to clarify lysates. The clarified lysate was then subjected to copper-based click chemistry analysis with tetramethyl-rhodamine azide (TAMRA) as a reporter group. Clarified lysate was mixed with 100 μM TAMRA (dissolved in DMSO) and 100 μM Tris[(1-benzyl-1H-1,2,3-triazol-4-yl) methyl] amine (TBTA), followed by the addition of 1 mM CuSO4 and 1 mM TCEP, dissolved in water. This mixture was allowed to react for 1 hour at 37 °C, mixing occasionally. The reaction was quenched by adding 6×SDS-PAGE loading buffer and the sample was separated on precast 4– 20% gel. S-acylated proteins, mainly GSDMD due to overexpression, were visualized by detecting fluorescence from the TAMRA channel using a ChemiDoc MP System.

### HeaTil click chemistry screen for zDHHCs

0.25 μg of wild-type human GSDMD-mCherry DNA was aliquoted to multiple 1.5 mL microfuge tubes. Human zDHHC constructs from the HeaTil screen (1–2 μg) (*32*) were individually added to each tube and mixed gently. Lipofectamine 2000 was added three times excess to the amount of total DNA and incubated for 20-30 minutes. This DNA-Lipofectamine mixture was then added to each well of a 6-well plate. GSDMD DNA, devoid of zDHHC, was also transfected as a control to check for background S-acylation. Cells were fed with 17-ODYA, harvested, and clicked as described above. Expression of yellow fluorescent protein (YFP)-tagged zDHHC and GSDMD proteins were detected by YFP fluorescence and western blot using anti-GSDMD antibody, respectively. Loading controls were determined by western blots developed with anti-GADPH antibody.

### Liposome leakage assay

Liposomes were prepared as previously described (*8, 44*). Briefly, phosphatidylcholine (1-palmitoyl-2-oleoyl-sn-glycero-3-phosphocholine, 25 mg/mL in chloroform; 80 μl), PE (1-palmitoyl-2-oleoyl-sn-glycero-3-phosphoethanolamine, 25 mg/mL in chloroform; 128 μl) and cardiolipin (CL, 1’,3’-bis(1,2-dioleoyl-sn-glycero-3-phospho)-sn-glycerol (sodium salt), 25 mg/mL in chloroform; 64 μl) were mixed and the solvent was evaporated under a stream of N_2_ gas. The lipid mixture was suspended in 1 ml buffer A (20 mM HEPES at pH 7.4, 150 mM NaCl, 50 mM sodium citrate, and 15 mM TbCl_3_) for 3 min. The suspension was pushed through 100 nm Whatman Nuclepore Track-Etched Membrane 30 times to obtain homogeneous liposomes. The filtered suspension was purified by size exclusion column (Superose 6, 10/300 GL) in buffer B (20 mM HEPES, 150 mM NaCl) to remove TbCl_3_ outside liposomes. Void fractions were pooled to produce a stock of PC/PE/CL liposomes (1.6 mM). Expression and purification of catalytically active caspase-1 were performed using the previously reported refolding method (*46*). In brief, two non-tagged p20 and p10 subunits were expressed in *E. coli* BL21 (DE3) bacteria in inclusion bodies individually. The p20/p10 complex was assembled by denaturation and refolding, which was further purified by HiTrap SP cation exchange chromatography (GE Healthcare Life Sciences).

The liposomes are diluted to 50 μM with buffer C (20 mM HEPES, 150 mM NaCl and 50 μM DPA) for use in a liposome leakage assay in which the leakage of Tb^3+^ from inside the liposomes was detected by an increase in fluorescence when Tb^3+^ was bound to dipicolinic acid (DPA) in buffer C. Human GSDMD (0.5 μM, WT or C191A mutant), bacterially produced or Expi293 expressed, was precleaved by 3C for 30 minutes or caspase-1 for 1 hour, and added to 384-well plates (Corning 3820) containing PC/PE/CL liposomes (50 μM liposome lipids). The fluorescence intensity of each well was measured at 545 nm using excitation at 276 nm immediately after mixing and followed for 2 hours using a SYNERGY microplate reader (Biotek).

### GSDMD and zDHHS Expression

zDHHS represents the catalytically inactive mutant of zDHHC in which the catalytic Cys is mutated to Ser. Expi293F cells were seeded at approximately 1×10^6^ cells/mL maintained in 50 mL Expi293 expression media and grown in an orbital shaker incubator at 37 °C, 105 rpm, and 5% CO_2_ until cells reached a density of 2×10^6^ cells/mL. Cells were transiently transfected with 50 μg of DNA: GSDMD (25μg) and zDHHS5/9/12/17/20 (25μg). GSDMD DNA devoid of zDHHS, and zDHHS DNA devoid of GSDMD were also transfected as controls to check for background resin binding. For transfection, DNA, purified in water, was mixed with polyethylenimine (PEI), at a ratio of 1:3 (DNA: PEI) in Expi293 expression media. After 30 minutes, the DNA-PEI mixture was added to the cells and grown in an orbital shaker incubator. 12 hours post-transfection, cells were fed with filtered 45% D-(+)-glucose and 10 mM sodium butyrate solution. Cells were left to incubate for an additional 48 hours. Cells were harvested with one wash of 1×PBS at 2500 × RPM for 30 minutes at 4 °C.

### GSDMD and zDHHS pulldowns (Co-immunoprecipitation)

GSDMD and zDHHS cell pellets were thawed on ice and resuspended in ice-cold cell lysis buffer (1×PBS pH 7.4, 1 mM TCEP, 1 mM EDTA, 10% glycerol, and 0.5% Triton X-100) supplemented with 100× protease inhibitor cocktail and benzonase nuclease. Cells were left to rotate for 90 minutes at 4°C and centrifuged at max speed for 10 minutes. The whole cell lysate was left to incubate with washed ANTI-FLAG® M2 Affinity Gel (Sigma-Aldrich A2220-4X25ML) at 4 °C for 2 hours on an end-over-end rotator. Samples were spun in a microcentrifuge at 500 × g for 2 minutes at 4 °C and the flow through was collected with a pipettor. Beads were washed 3 times with ice-cold Wash Buffer 1 (1×PBS pH 7.4, 1 mM TCEP, 1 mM EDTA, 5% glycerol, and 0.5% Triton X-100). Beads were washed a final time with ice-cold Wash Buffer 2 (1×PBS pH 7.4, 1 mM EDTA, 5% glycerol, and 0.5% Triton X-100). Samples were eluted by incubating at room temperature on a shaking rack for 5 minutes with elution buffer (1×PBS pH 7.4, 1 mM EDTA, 5% glycerol, 0.5% Triton X-100, and 150 ng/µL 3X FLAG peptide) and spun at 500 × g in a microcentrifuge for 1 min. The supernatant was carefully collected with a pipettor and transferred to a clean microcentrifuge tube appropriately labeled with the sample and fraction number. Elution was repeated four more times. The fractions were then separated by SDS-PAGE and analyzed by Western blotting for GSDMD (anti-FLAG) and zDHHC5 or zDHHC9 (antibody).

### Quantification and statistical analysis

Statistical significance was calculated by using GraphPad Prism 6.01 (unpaired Student’s test, https://www.graphpad.com/scientific-software/prism/). The number of independent experiments of duplicates, the statistical significance, and the statistical test used to determine the significance are indicated in each figure or figure legend or method section where quantification is reported.

## Supporting information

Suppemental figures

## Acknowledgements

We would like to thank Daniel Bachovchin for GSDMD KO THP-1 cells, and the Harvard Medical School Nikon Imaging Center for help with fluorescence microscopy. The graphical diagram was created with Biorender.com. This work was supported by US National Institutes of Health grants R01AI139914 (H.W. and J.L.), DP1HD087988 (H.W.), R01AI124491 (H.W.), R01CA240955 (J.L.). C.W. received postdoctoral fellowship from NIGMS. P.F. received postdoctoral fellowship from the Cancer Research Institute. Y.D. received postdoctoral fellowship from the Charles A. King Trust.

