## Supplementary material for "ROS-dependent palmitoylation is an obligate licensing modification for GSDMD pore formation": Suppemental figures: Supplemental Materials.pdf

The file includes:
Figs. S1 to S7

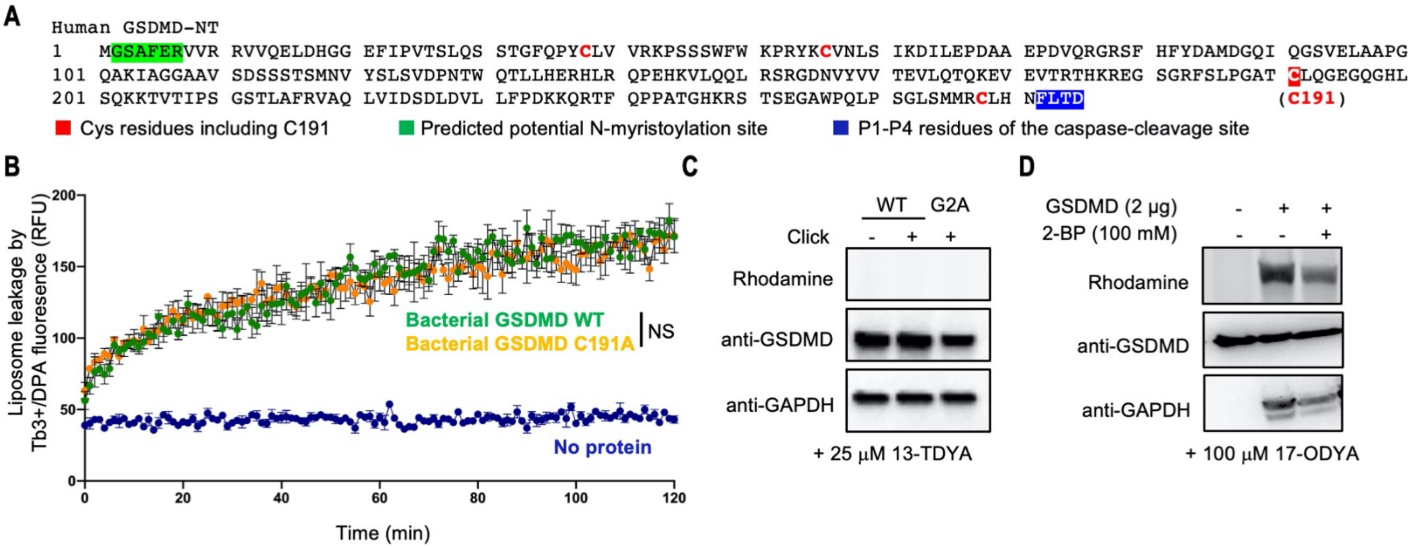


**Fig. S1. GSDMD is highly palmitoylated but not myristoylated.** (**A**) Protein sequence of human GSDMD N-terminal domain (GSDMD-NT). Certain residues are marked by colors. (**B**) Liposome leakage by precleaved bacterium-expressed WT GSDMD in comparison to precleaved C191A mutant GSDMD revealed similar activity of WT and C191A. Results were obtained from 3 independent experiments (n = 3) and expressed as mean ± SEM. Two-way ANOVA was used to assess the statistics with NS (non-significant) for p>0.05. (**C**) Click chemistry for protein N-myristoylation did not detect N-myristoylation of GSDMD. **(D)** GSDMD-FL palmitoylation detected by click chemistry, and its inhibition by the general palmitoylation inhibitor 2-BP.

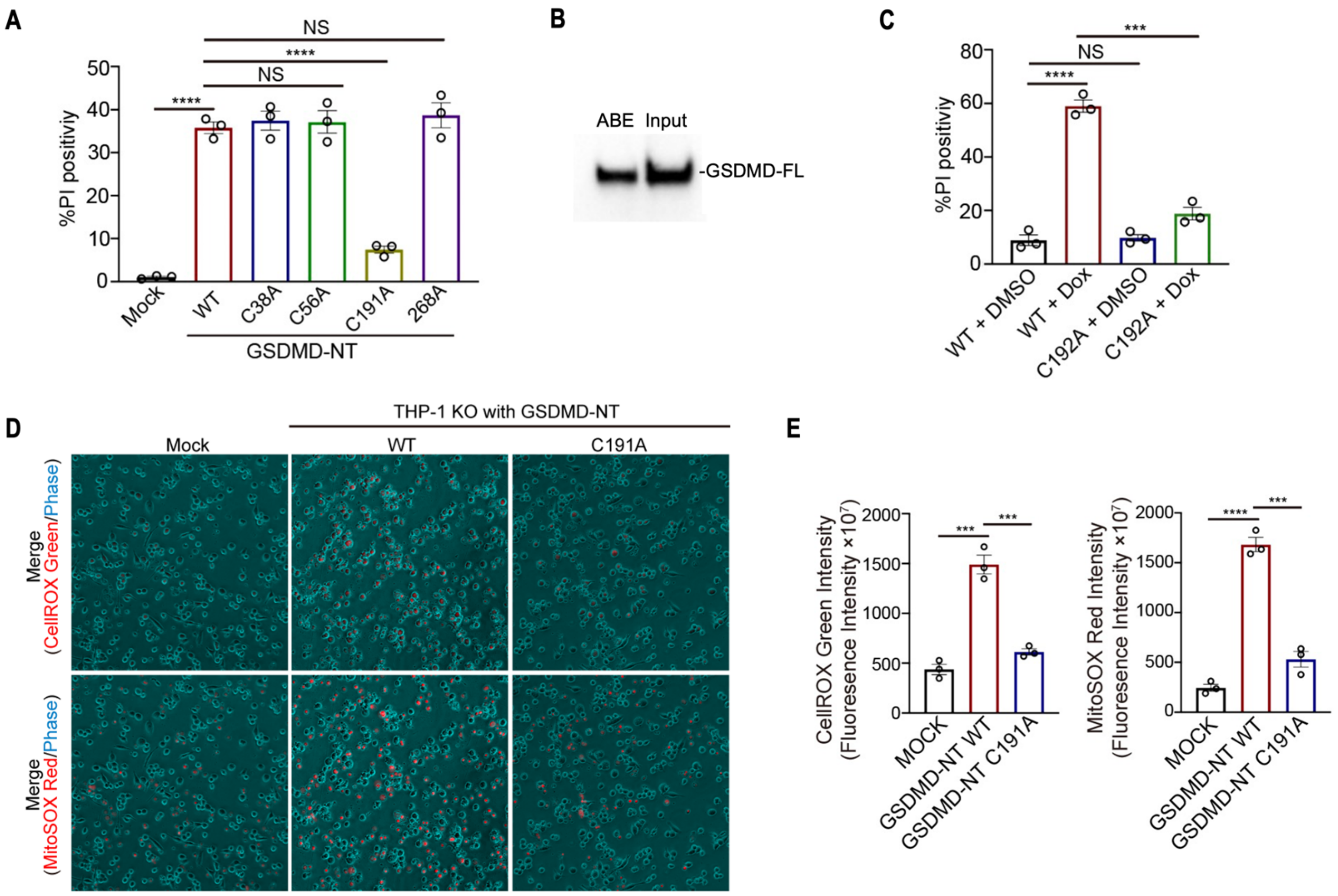


**Fig. S2. GSDMD is specifically palmitoylated at Cys191/192 (human/mouse) and defectiveness in palmitoylation compromises liposome leakage *in vitro*, and GSDMD membrane localization and pyroptosis in cells.** (**A**). PI positivity of HEK293T cells overexpressing WT or C191A GSDMD-NT, showing impairment of the C191A mutant in inducing pyroptosis. (**B**) ABE on the Expi293 cell-expressed GSDMD-FL sample used for liposome leakage, showing the palmitoylation of GSDMD-FL and the percentage of palmitoylation at ~27%. The input lane was 2-fold diluted. (**C**) PI positivity GSDMD KO iBMDMs reconstituted with Dox-inducible WT or C192A mouse GSDMD-NT, showing impairment of the C192A mutant in inducing pyroptosis. (**D**) Measurement of cellular oxidative stress by CellROX Green and mitochondrial ROS by MitoSOX red in GSDMD KO THP-1 cells reconstituted with WT or C191A GSDMD-NT, showing the activation of cellular oxidative stress and mitochondrial ROS of WT GSDMD-NT, not C191A. **(E)** Quantification of fluorescence intensity of CellROX Green and MitoSOX red in (D). All results were obtained from at least 3 independent experiments. Error bars represent SEM. Statistics used Student's t-tests with *** for $p<0.001$, **** for $p<0.0001$, and NS (non-significant) for $p>0.05$.

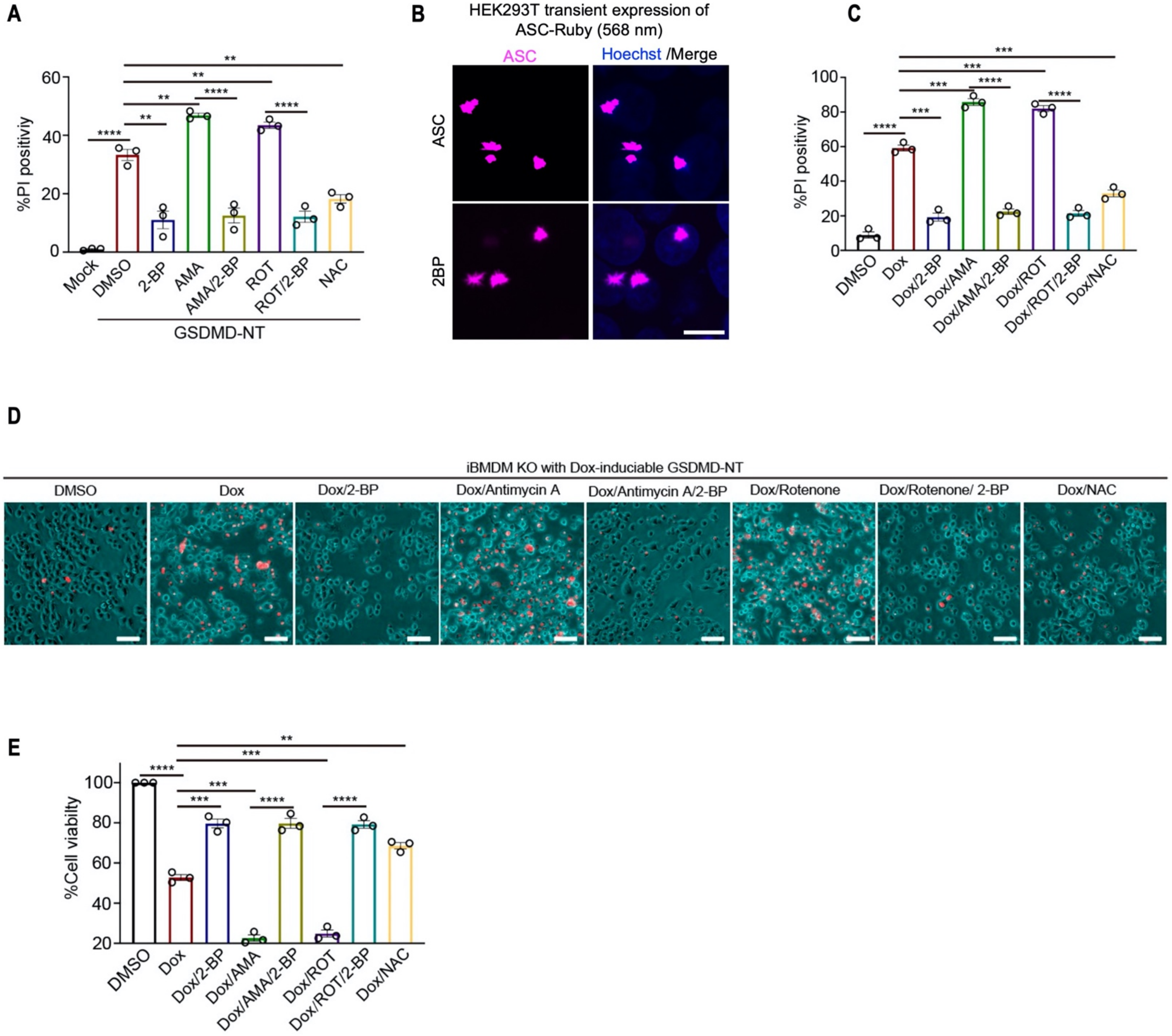


**Fig. S3. Regulation of GSDMD palmitoylation and pyroptosis by ROS modulators in HEK293T cells and iBMDMs.** (**A**) PI positivity of HEK293T cells overexpressing GSDMD-NT and treated with ROS modulators, and 2-BP or not, showing enhanced pyroptosis by Rot and AMA, and decreased pyroptosis by NAC. (**B**) Localization analysis of ASC after expression of ASC-mRuby in HEK293T, showing no difference in ASC cellular localization with or without 2-BP treatment. (**C**) PI positivity of GSDMD KO iBMDMs reconstituted with Dox-inducible GSDMD-NT, showing enhanced pyroptosis by Rot and AMA and its inhibition by 2-BP, and decreased pyroptosis by NAC. (**D**) PI and phase merged images of GSDMD KO iBMDMs reconstituted with Dox-inducible GSDMD-NT treated with ROS modulators. (**E**) cell viability of GSDMD KO iBMDMs reconstituted with Dox-inducible GSDMD-NT, showing enhanced pyroptosis by Rot and AMA and its inhibition by 2-BP, and decreased pyroptosis by NAC. All results were obtained from at least 3 independent experiments. Scale bars represent 25 μm (B)

and 50 µm (D). Error bars represent SEM. Statistics used Student's t-tests, with ** for $p<0.01$, *** for $p<0.001$, and **** for $p<0.0001$.

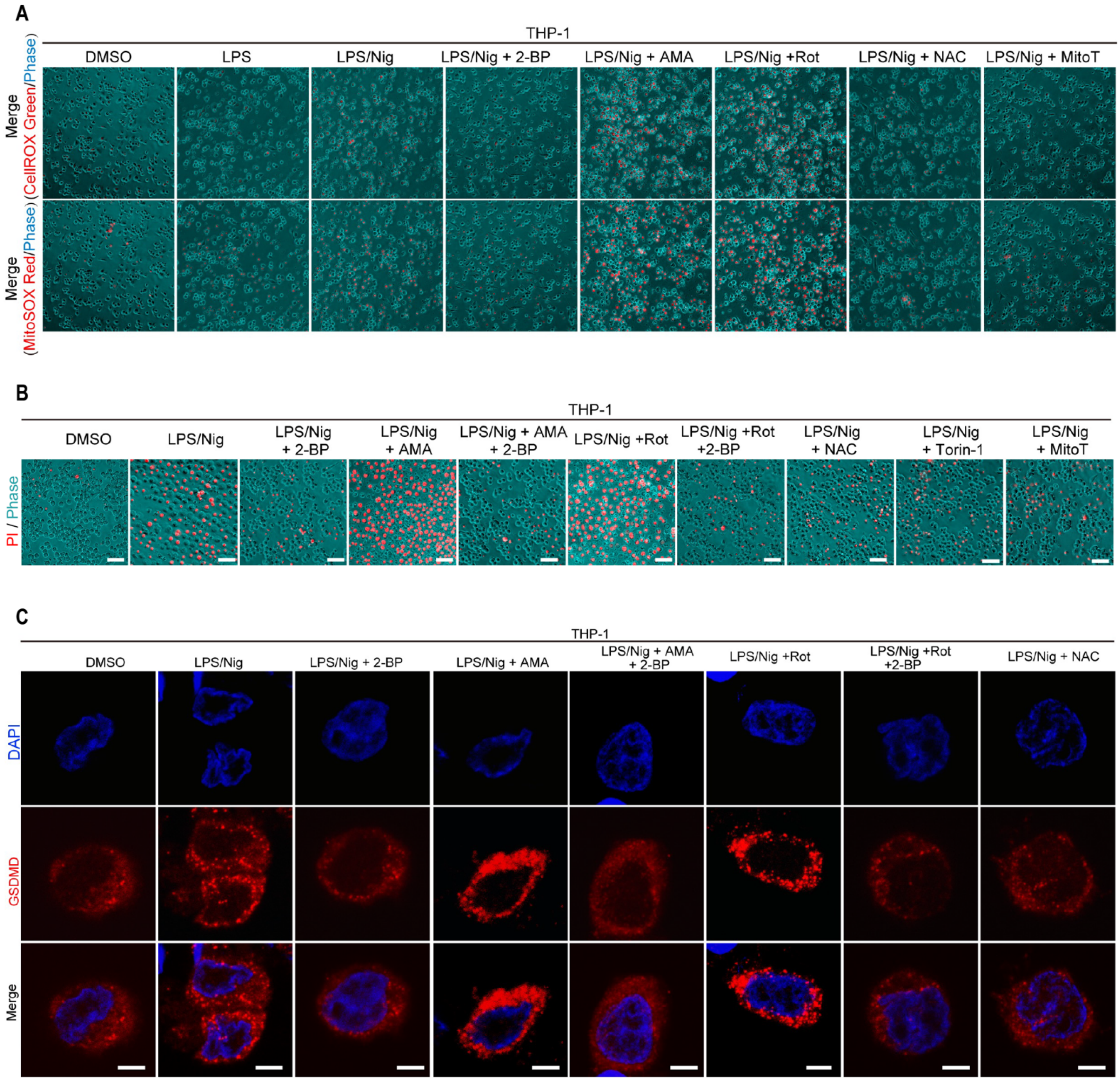


**Fig. S4. Regulation of GSDMD palmitoylation and pyroptosis by ROS modulators in THP-1 cells.** (**A, B**) Merged images of CellROX Green and phase, MitoSOX Red and phase (A) and merged images of PI and phase (B) in THP-1 cells treated with ROS modulators and LPS plus nigericin. (**C**) Anti-GSDMD immunofluorescence imaging of THP-1 WT cells treated with LPS plus nigericin and ROS modulators, showing increased cell membrane localization by Rot and AMA and its inhibition by 2-BP, as well as the diffuse cytoplasmic localization upon treatment by NAC, Torin-1, or MitoT.

DAPI (blue) and PI (Red) stain nuclei. Results were obtained from 3 independent experiments. Scale bars represent 50 µm (B), and 5 µm (C).

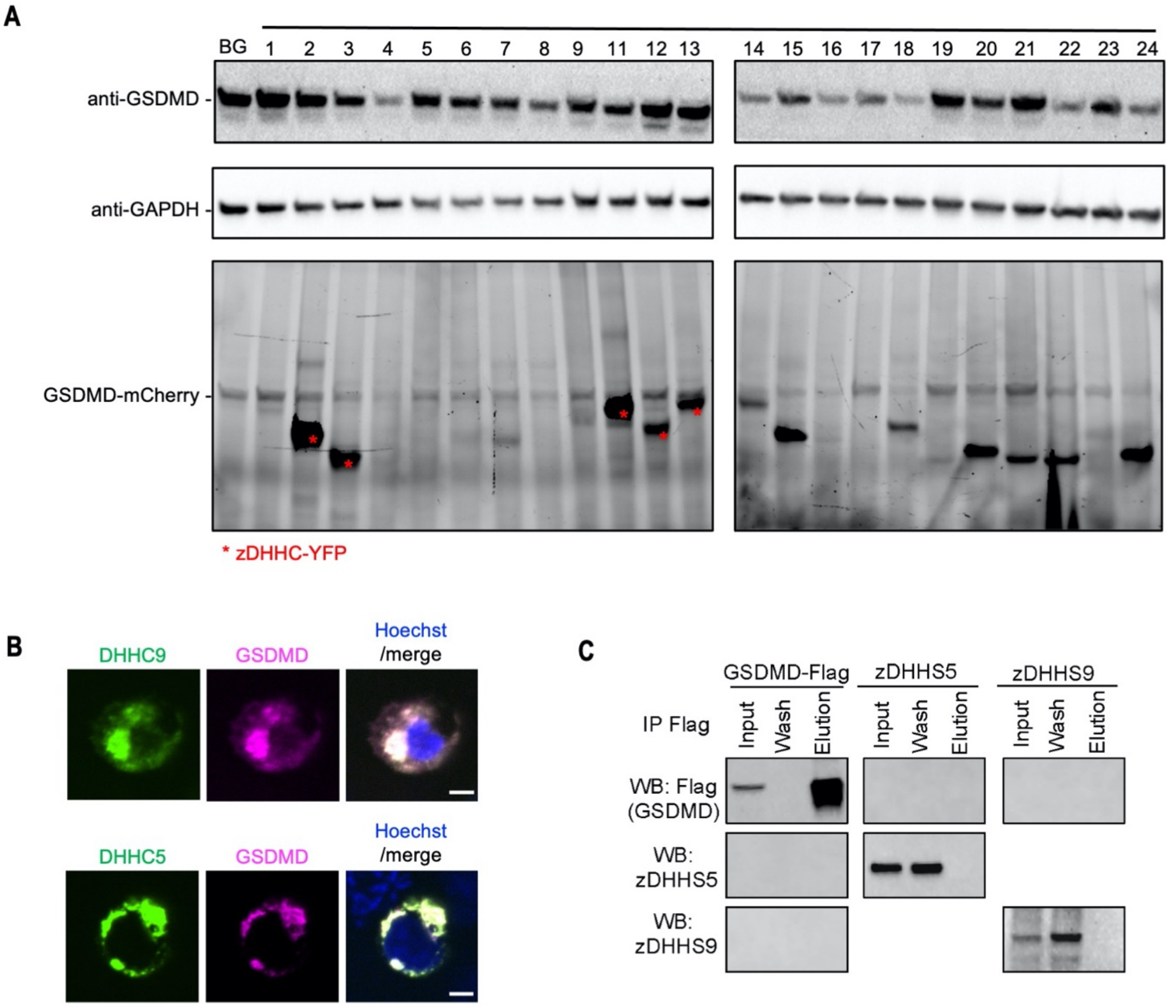


**Fig. S5. Identification of zDHHC5 and zDHHC9 as the main palmitoyltransferases for GSDMD palmitoylation.** (**A**) Control panels for click chemistry screen by rhodamine labeling in 293T cells co-expressing the 23 human zDHHCs with GSDMD to identify zDHHCs that enhanced GSDMD palmitoylation. GAPDH was used as a loading control. (**B**) Co-localization imaging analysis of zDHHC5-YFP and zDHHC9-YFP with GSDMD-mCherry upon co-expression in HEK293T cells. Hoechst (blue) stained nuclei. The cap-like structures appear to be intact Golgi. (**C**) Anti-FLAG pulldown control. zDHHS5 and zDHHS9 were not pulled down without co-expression with the GSDMD-FLAG bait, and GSDMD-FLAG cannot pulldown zDHHS5 and zDHHS9 without the expression of zDHHS5 and zDHHS9. All results were obtained from at least 3 independent experiments. Scale bars represent 5 μm (B).

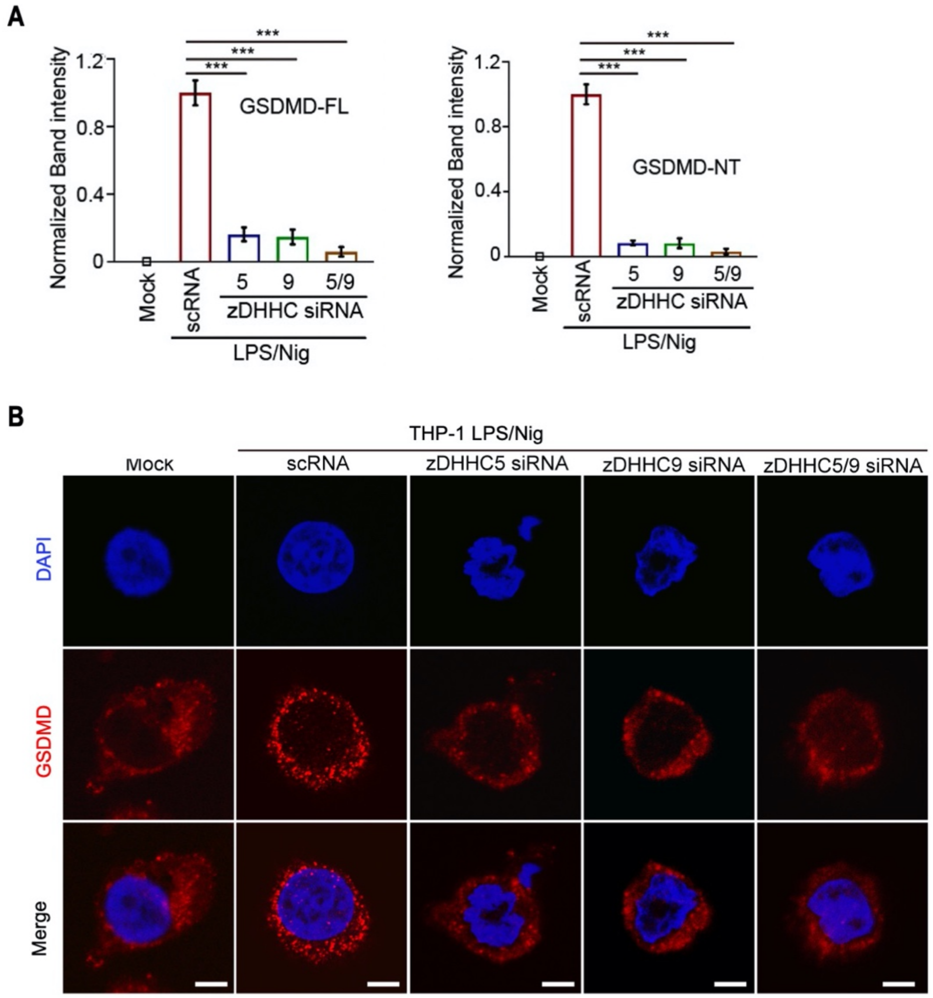


**Fig. S6. Identification of zDHHC5 and zDHHC9 as the main palmitoyltransferases for GSDMD palmitoylation.** (**A**) Quantification of band intensity of GSDMD-FL and GSDMD-NT palmitoylation detected by ABE in THP-1 cells upon siRNA knockdown of zDHHC5, zDHHC9, or both and treatment with LPS plus nigericin, showing that these knockdowns compromised GSDMD palmitoylation. **(B)** Anti-GSDMD immunofluorescence imaging of THP-1 cells with siRNA knockdowns of zDHHC5, zDHHC9, or both and treatment with LPS plus nigericin. Only cells treated with scRNA showed strong cell surface staining. zDHHC5 siRNA, zDHHC9 siRNA, or both showed the inhibition of GSDMD cell surface localization. DAPI (blue) stained nuclei. Scale bars represent 5 μm. All results were obtained from 3 independent experiments. Error bars represent SEM. Statistics used Student’s t-tests, with *** for $p<0.001$.

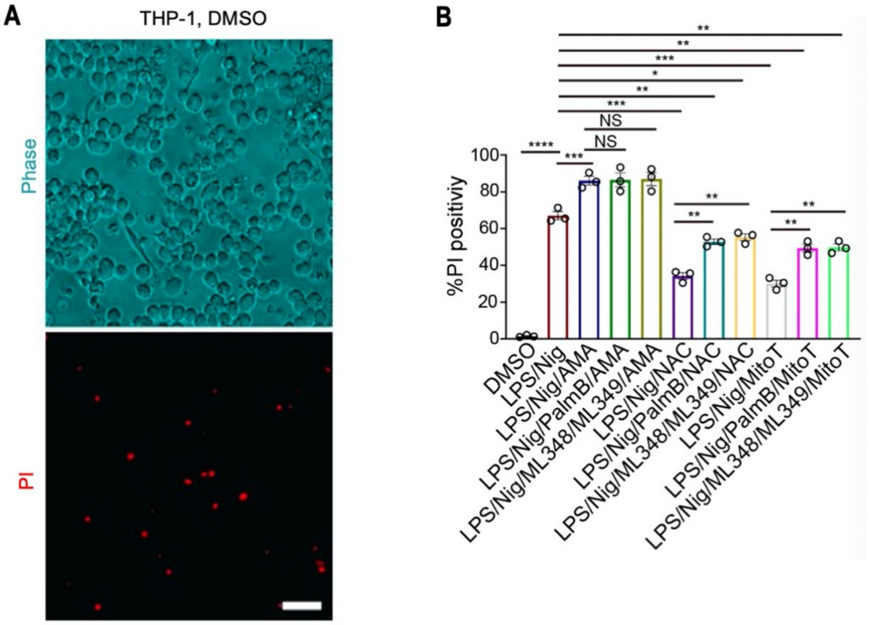


**Fig. S7. Mechanism of ROS-regulated GSDMD palmitoylation.** (**A**) Negative control of Fig. 4C**.** (**B**) PI quantification of THP-1 cells with the treatment of LPS/Nig and ROS modulators, with or without treatment of depalmitoylase inhibitors, PalmB (general inhibitor), ML348 (APT1 inhibitor) and ML349 (APT2 inhibitor). Results were obtained from 3 independent experiments. Error bars represent SEM. Statistics used Student's t-tests, with ** for p<0.01, *** for p<0.001, **** for p<0.0001, and NS (non-significant) for p>0.05.
